# Engineering a material-genetic interface as safety switch for embedded therapeutic cells

**DOI:** 10.1101/2023.04.03.535359

**Authors:** Carolina Jerez-Longres, Marieta Gómez-Matos, Jan Becker, Maximilian Hörner, Franz-Georg Wieland, Jens Timmer, Wilfried Weber

**Affiliations:** INM – Leibniz Institute for New Materials, Campus D2 2, 66123 Saarbrücken, Germany; Department of Materials Science and Materials Engineering, Saarland University, 66123 Saarbrücken, Germany; Signalling Research Centres BIOSS and CIBSS, University of Freiburg, Schänzlestrasse 18, 79104 Freiburg, Germany; Faculty of Biology, University of Freiburg, Schänzlestrasse 1, 79104 Freiburg, Germany; SGBM - Spemann Graduate School of Biology and Medicine, University of Freiburg, Albertstrasse 19a, 79104 Freiburg, Germany; Institute of Physics, University of Freiburg, Hermann-Herder-Strasse 3, 79104 Freiburg, Germany; Freiburg Center for Data Analysis and Modelling (FDM), University of Freiburg, Ernst-Zermelo-Strasse 1, 79104 Freiburg, Germany

## Abstract

Encapsulated cell-based therapies involve the use of genetically-modified cells embedded in a material in order to produce a therapeutic agent in a specific location in the patient’s body. This approach has shown great potential in animal model systems for treating diseases such as type I diabetes or cancer, with selected approaches having been tested in clinical trials. Despite the promise shown by encapsulated cell therapy, though, there are safety concerns yet to be addressed, such as the escape of the engineered cells from the encapsulation material and the resulting production of therapeutic agents at uncontrolled sites of the body. For that reason, there is great interest in the implementation of safety switches that protect from those side effects. Here, we develop a material-genetic interface as safety switch for engineered mammalian cells embedded into hydrogels. Our switch allows the therapeutic cells to sense whether they are embedded in the hydrogel by means of a synthetic receptor and signaling cascade that link transgene expression to the presence of an intact embedding material. The system design is highly modular, allowing its flexible adaptation to other cell types and embedding materials. This autonomously acting switch constitutes an advantage over previously described safety switches, which rely on user-triggered signals to modulate activity or survival of the implanted cells. We envision that the concept developed here will advance the safety of cell therapies and facilitate their translation to clinical evaluation.

**Graphical abstract:** 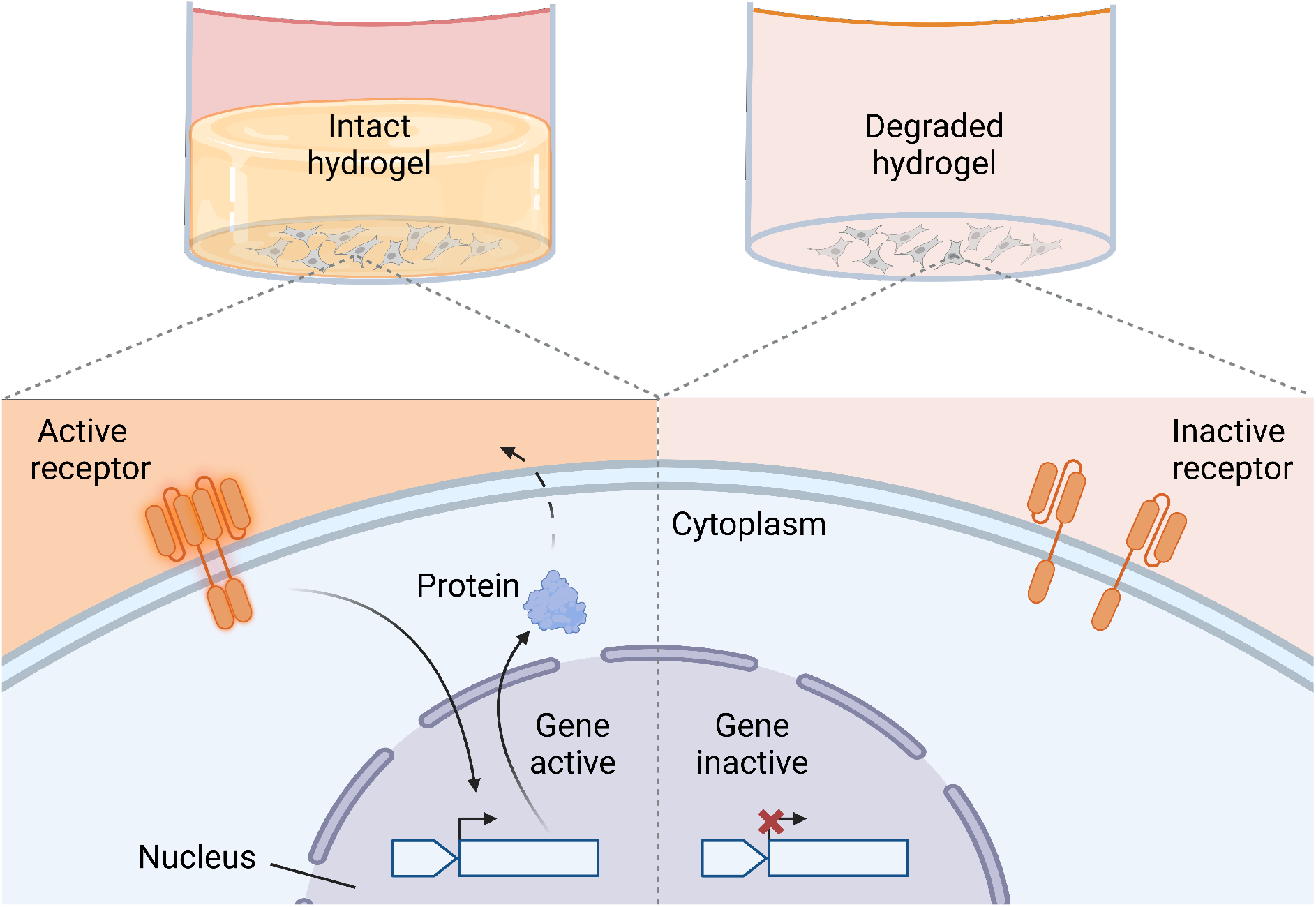

## 1 Introduction

Cell-based therapies employ engineered mammalian cells for the production of a therapeutic agent in the patient’s body, bypassing the need for frequent applications of exogenously added drugs such as recombinant proteins. Typically, cells produce the therapeutic proteins either in response to user-triggered external cues or to specific biomarkers in order to seamlessly adapt therapeutics production to changing patient needs^1^. For instance, human embryonic kidney cells have been engineered as pancreatic islet cell mimics to sense glucose levels and produce insulin for treatment of type I diabetes^2^. Likewise, allogenic mesenchymal stem cells, that have a tropism for tumors, can be used to deliver drugs at the disease site^3,4^. Several cell-based therapeutics have already made it to the clinic or are undergoing clinical studies, while numerous others are under pre-clinical investigation^5,6^.

Despite the promising potential of cell-based therapies to treat major and chronic diseases, and the translation to the clinics of some of them, important challenges remain that need to be addressed. One major safety concern is the uncontrolled distribution of cells in the body and the resulting production of recombinant proteins at uncontrolled body sites. This issue is commonly overcome by cell encapsulation into polymeric materials which provides a means of implanting and maintaining the therapeutic cells at the desired location, while simultaneously ensuring the transport of nutrients into the implant and of therapeutic agents outside of it^7^. In addition, it allows for lower doses and potentially less side effects^8–10^. The idea was first successfully applied in preclinical studies with encapsulated pancreatic islet cells to control diabetes^11^. Since then, research with encapsulated mammalian cells has been carried out in other fields, such as degenerative bone disease, hepatic disease, neurological disease, cancer^12,13^ and urate homeostasis, to name a few^8,14,15^. Although the gap to regulatory approval of encapsulated cell therapy has not been closed yet^9,16^, several clinical trials are underway, for example with encapsulated genetically modified cells which release neurotrophic factors for neurodegenerative diseases^17,18^. Additionally, other application areas are opening up in encapsulated cell therapy, such as the use of encapsulated cells for ocular drug delivery^19^ or encapsulation of stem cells for regenerative medicine^20–22^.

An important consideration to make when designing encapsulated cell therapeutics is the choice of encapsulation material. An ideal encapsulation matrix should have material properties similar to those of the extracellular matrix (ECM). This is the case for hydrogels, whose high water content and viscoelastic properties make them highly cell-compatible^23^. Secondly, the material should ideally be made of clinically approved molecules and have a good safety profile. Hydrogels typically used for cell encapsulation can be of biological or synthetic origin^8,10^. Among the former, the biomacromolecules of choice are usually polysaccharides, such as alginate, agarose and hyaluronic acid, and proteins, among which collagen stands out as the main component of the ECM, alongside others such as fibrin and elastin^9,16^. Synthetic polymers, on the other hand, have the advantage of easier processing and better mechanical stability^24^. The most widely used among these is polyethylene-glycol (PEG), being FDA-approved since 1995^25^ and having been used for several drug formulations since^26^. Due to its excellent biocompatibility and physicochemical properties (such as low protein adsorption, which results in less immunogenicity^27^), PEG is highly compliant with the requirements for therapeutic cell encapsulation.

Despite advances in cell encapsulation, critical safety concerns remain. Importantly, escape of the engineered cells from the encapsulation material could occur, either due to degradation of the material^28,29^ or due to cell proliferation and outgrowth from the matrix^29–32^. Such escape would lead to production of the therapeutic protein at uncontrolled sites of the body, which would induce off-target toxicity, especially in cases in which local administration is important, such as localized tumor therapy^12,13^. Translating such approaches to the clinic therefore requires safety switches that prevent uncontrolled protein production once cells escape the embedding material.

Various safety switches for cell therapy have been developed and a few are deployed in clinical studies, as is the case with *Herpes simplex* thymidine kinase^33,34^ or inducible caspase^19,35^ suicide switches. Other approaches use engineered cells that are dependent on an exogenously administered metabolite^36^. All these switches are actuated by externally applied molecules and result in the killing of all implanted cells. Thus, the safety switch can only be actuated once after uncontrolled cell release and off-target toxicities are detected. However, an optimal switch would be able to autonomously detect whether a cell escaped the embedding material and then selectively inactivate therapeutic protein production only in these cells. Such a switch would allow for longer treatment durations while, at the same time, preventing side effects caused by engineered cells being active outside the implant. In this study, we develop a material-genetic interface that directly links transgene expression to the presence of an embedding hydrogel to be used as safety switch to shut off protein production once cells escape the material.

In order to build such a switch, the cells have to be engineered to sense their environment. Gene networks allow to implement such sensing and responding capabilities into mammalian cells^37^. To detect a certain stimulus, chimeric synthetic receptors have been engineered that expand the sensing capabilities of cells to molecules that have no natural receptors^1^. Recently, a generic concept for the design of synthetic receptors has been developed. The GEMS (Generalized Extracellular Molecule Sensor) platform^38^ comprises a modular chimeric receptor, activated by dimerization, and whose intracellular domain can be selected to signal through different endogenous pathways, resulting in the modulation of transgene expression.

In this study, we engineered the GEMS platform to detect clustered fluorescein molecules, a clinically licensed compound routinely used as contrast agent in ophthalmology. Embedding cells in a PEG-based fluorescein-functionalized hydrogel thus induces transgene expression whereas the degradation of the material or cultivation of cells outside the material results in de-activation of the synthetic signaling cascade and thus the downregulation of target gene expression. We characterize the autonomous functionality of this safety switch and demonstrate a strict correlation between transgene expression levels and the intactness of the embedding material. We anticipate that the underlying concept and the modular nature of the switch will foster advances in promoting the safety of cell therapies towards clinical translation.

## 2 Material and methods

### 2.1 Construction of expression vectors

To engineer the synthetic receptor to bind fluorescein (pCJL503), the sequence coding for the fluorescein-specific single chain variable fragment (scFv) FITC-E2^39^ was amplified from plasmid pOT021 (unpublished) and inserted by Gibson assembly into pLeo669^38^ (kindly provided by Martin Fussenegger) to replace the SunTag-specific scFv. For flow cytometry readouts, a fluorescent reporter (pCJL510) was generated by amplifying the mScarlet gene from pMH703 (unpublished) and cloning it into pMF111^40^ under the control of the tetracycline-responsive promoter. All plasmids generated in this study are listed in the Supplementary Material together with the corresponding DNA sequences.

### 2.2 Conjugation of fluorescein to bovine serum albumin (BSA)

BSA (Carl Roth, Karlsruhe, Germany, cat. no. T844.2) was dissolved in phosphate buffered saline (PBS; 137 mM NaCl, 8 mM Na_2_HPO_4_, 2.7 mM KCl, 1.5 mM KH_2_PO_4_, pH 7.2) at a concentration of 10 mg/mL and fluorescein isothiocyanate (FITC; Sigma-Aldrich, St. Louis, MO, cat. no. F7250) was added in a 25x molar excess from a 10 mg/mL solution in dimethylsulfoxide (DMSO). After incubation for 1 h at room temperature (RT), the uncoupled FITC was removed by two dialysis steps (10 kDa molecular weight cutoff, MWCO) against PBS at RT for 2 h and one final step at 4 °C over night. BSA has 60 primary amines, which are potential conjugation sites for FITC.

### 2.3 3C protease (3CP) production and purification

*E. coli* BL21(DE3)-pLysS cells (Thermo Fisher Scientific, Waltham, MA, cat. no. C602003) were transformed with plasmid pHJW257 (unpublished), coding for strep-tagged 3CP, and selected by growth in Luria/Miller broth (LB) supplemented with ampicillin (100 μg/mL) and chloramphenicol (36 μg/mL). Bacteria were grown in LB medium in flasks at 37 °C while shaking to an OD_600_ of 0.9 prior to induction with 1 mM isopropyl β-D-1-thiogalactopyranoside (IPTG; Carl Roth, cat. no. 2316.5) and a protein production period of 5 h at 37 °C. Subsequently, the bacteria were harvested by centrifugation and the cells were resuspended in column buffer (100 mM Tris/HCl, 150 mM NaCl, pH 8.0, 35 mL per 1 L of initial culture), shock-frozen in liquid nitrogen and stored at −80 °C until purification. For affinity chromatography purification, resuspended pellets were lysed by ultrasonication (Sonoplus HD, Bandelin, Berlin, Germany). The lysates were clarified by centrifugation at 30 000 g for 30 min and the supernatant was loaded onto a gravity-flow column containing StrepTactin XT 4Flow resin (IBA Lifesciences, Göttingen, Germany, cat. no. 2-5030-002; 1.5 mL StrepTactin beads per 1 L bacterial culture) previously equilibrated with column buffer. The 1.5 mL column was washed with 15 mL column buffer, after which the protein was eluted in 7 fractions of 1 mL each with elution buffer (100 mM Tris/HCl, 150 mM NaCl, 50 mM biotin, pH 8.0). The fractions with the highest absorbance at 280 nm were pooled and β-mercaptoethanol was added to a final concentration of 10 mM. The eluate was concentrated using a Vivaspin 5k-MWCO concentrator (Sartorius AG, Göttingen, Germany, cat. no. VS04T11), and the final concentration was determined by Bradford assay. Finally, 10% (v/v) glycerol was added and the protein was shock-frozen in single-use aliquots in liquid nitrogen and stored at −80 °C.

### 2.4 Cell culture and transient transfection

Human embryonic kidney cells (HEK-293T, DSMZ, Braunschweig, Germany, cat. no. ACC 635) were cultivated in Dulbecco’s modified Eagle’s complete medium (DMEM; Pan Biotech, Aidenbach, Germany, cat. no. P04-03550) supplemented with 10% (v/v) fetal bovine serum (FBS; PAN Biotech, cat. no. P30-3602) and 1% (v/v) streptomycin/penicillin solution (PAN Biotech, cat. no. P06-07100) in a humidified atmosphere containing 5% CO_2_ at 37 °C.

For transfection, tissue culture-treated 96-well plates (Corning, Corning, NY, cat. no. 3599) were coated with collagen (Life Technologies, Carlsbad, CA, cat. no. A1048301) at 50 μg/mL in 25 mM acetic acid by incubating at RT for 2 h and subsequent washing with PBS. Plates were stored at 4 °C until use. Cells were seeded 16 h before transfection on the pre-coated plates at a density of 1 x 10^4^ cells/well (for reversibility experiments) or 3 x 10^4^ cells/well (for dose-response and hydrogel experiments) in 100 μL DMEM complete medium per well. Cells were transfected via polyethyleneimine (PEI; Polysciences, Warrington, PA, cat. no. 23966-1). The protocol was adapted from Longo *et al.^41^*, using 126 ng DNA and 420 ng PEI in 16.8 μL OptiMEM medium (Life Technologies, Carlsbad, CA, cat. no. 22600-134) per well of a 96-well plate. Plasmid ratios (in μg) were as follows: pCJL503:pMF111:MKp37 = 3:1:1 for secreted alkaline phosphatase (SEAP) assay readout and pCJL503:pCJL510:MKp37:pOK045 = 57:19:19:5 for flow cytometry readout.

### 2.5 Receptor activation with BSA-Fluorescein

HEK-293T cells were seeded and transfected on 96-well plates as described above. 5 h after transfection, the cell culture medium containing the transfection mix was exchanged by DMEM complete medium containing BSA-fluorescein at the indicated concentrations. For negative controls with free fluorescein, DMEM complete medium was supplemented with 200 μM sodium fluorescein (Sigma-Aldrich, cat. no. F6377) from a 25 mM stock solution in water. After addition of the compounds, the cells were incubated for 20 h before analysis.

### 2.6 Hydrogel synthesis and dissolution

40 kDa 8-arm PEG-vinylsulfone (PEG-VS) was obtained from NOF Europe (Frankfurt, Germany, cat. no. Sunbright HGEO-400V) and dissolved in triethanolamine (TEA) buffer (300 mM triethanolamine-HCl, pH 8.0) at a concentration of 25% (w/v). Fluorescein-labelled and crosslinker peptides were obtained from Genscript (Piscataway, NJ) in lyophilized form. Before each experiment, the peptides were dissolved in TEA buffer at 10 mM concentration (fluorescein-labelled peptide) and water at 15 mM (crosslinker peptide). The sequences of the designed peptides are listed in **Table S2** in the Supplementary Material.

HEK-293T were seeded and transfected in 96-well plates as described above. 5 h after transfection, hydrogels were synthesized as follows. Per well, the following reaction mix was prepared: 6 μL PEG- VS (resulting in a final concentration of 5% (w/v)) and 0-6 μL FITC-labelled peptide (corresponding to a concentration of 0-2 mM in the final hydrogel), filled up with TEA buffer to a final gel volume of 30 μL. The mix was incubated for 10 min at 37 °C. Subsequently, 10 μL crosslinker peptide (corresponding to a 4:1 peptide:PEG-VS molar ratio) were added to the mix, the cell culture medium was removed and the 30 μL-hydrogel mix was pipetted onto the cells. After polymerization for 3 min at 37 °C, the gels were washed 5 times for 5 min each with 250 μL DMEM complete medium (added on top of the gel). After the last washing step, 90 μL DMEM complete medium were added. For hydrogel dissolution, 10 μL 3CP (3 mg/ml in 100 mM Tris/HCl, 150 mM NaCl, 50 mM biotin, pH 8.0) were added to the medium. For negative controls, 10 μL 3CP buffer were added. Subsequently, the cells were incubated for 20 h before analysis.

### 2.7 SEAP activity assays

For SEAP analysis, cell culture medium was taken, heat-inactivated at 65 °C for 45 min and centrifuged for 1 min at 1250 g. Subsequently, 80 μL supernatant were mixed with 100 μL 2x SEAP buffer (20 mM homoarginine-HCl, 1 mM MgCl_2_, 21% (v/v) diethanolamine, pH 9.8) and 20 μL 20 mM para-nitrophenylphosphate (pNPP) in water, per well of a 96 well plate (Carl Roth, cat. no. 9293.1). The absorbance at 405 nm was measured every 1 min for at least 30 min and up to 2 h using a Synergy 4 microplate reader (Biotek, Winooski, VT). SEAP activities were calculated as a function of the increase in absorbance at 405 nm over time^42^.

### 2.8 Flow cytometry analysis

For flow cytometry readout of hydrogel samples, a fluorescent mScarlet (pCJL510) reporter was used. For internal normalization, a constitutive expression vector (pOK045, unpublished) for the blue fluorescent protein (BFP) was used. The samples were prepared as follows. For hydrogel samples without 3CP, 3CP was added 2 h before flow cytometry so as to dissolve the hydrogels.

Subsequently, the medium was removed from all samples, cells were detached with trypsin by 1 min incubation at 37 °C, and cells were resuspended in 250 μL PBS with 10% FBS and kept on ice until analysis. The samples were analyzed on an Attune NxT flow cytometer (Thermo Fisher Scientific) equipped with a 405 nm laser (for BFP) and a 561 nm laser (for mScarlet).

For receptor surface expression analysis, cell culture medium was removed and cells were incubated with trypsin for 1 min at 37 °C to detach the cells. These were resuspended in 250 μL PBS with 10% FBS and kept on ice until analysis on an Attune NxT flow cytometer using a 488 nm laser to detect fluorescein.

### 2.9 Rheological characterization of hydrogels

Hydrogels (50 μL volume, 5% (w/v) PEG, fluorescein-labeled peptide:VS = 1:10, see section 2.6) were prepared between glass slides previously treated with Sigmacote (Sigma-Aldrich, cat. no. SL2) (distance 1 mm). After polymerization for 3 min at RT, hydrogels were transferred to PBS and stored at 4 °C over night for swelling, and up to 1 week before analysis. Oscillatory shear measurements were carried out on a MCR301 rheometer (Anton Paar, Graz, Austria) equipped with parallel plates at 37 °C. The hydrogels were placed on the lower plate surrounded by a 3D-printed poly-lactic acid ring, with an inner diameter of 11 mm, to keep the gel and surrounding buffer in place, and compressed with an upper plate (PP08, Anton Paar, 0.05 N normal force). The storage and loss moduli were recorded over a time period of about 2 h at a frequency of 1 Hz and with a 0.5 % deformation (for frequency and amplitude sweeps, see **Fig. S3**). PBS with 3CP (0.3 mg/mL) and 10 mM β-mercaptoethanol (to preserve 3CP activity) was added at 20 min intervals.

### 2.10 Software

Flow cytometry data analysis, and Figures S1 and S2, were made using FlowJo (version 10.8.1, DB, Franklin Lakes, NJ). Rheology analysis was performed using Rheoplus (version 3.41, Anton Paar). Figures 3d and S3 were created using OriginPro (version 2020, OriginLab, Northampton, MA). Figures 1, 2a, 3a-b, 4a and the graphical abstract were created using BioRender.com.

**Figure 1.**
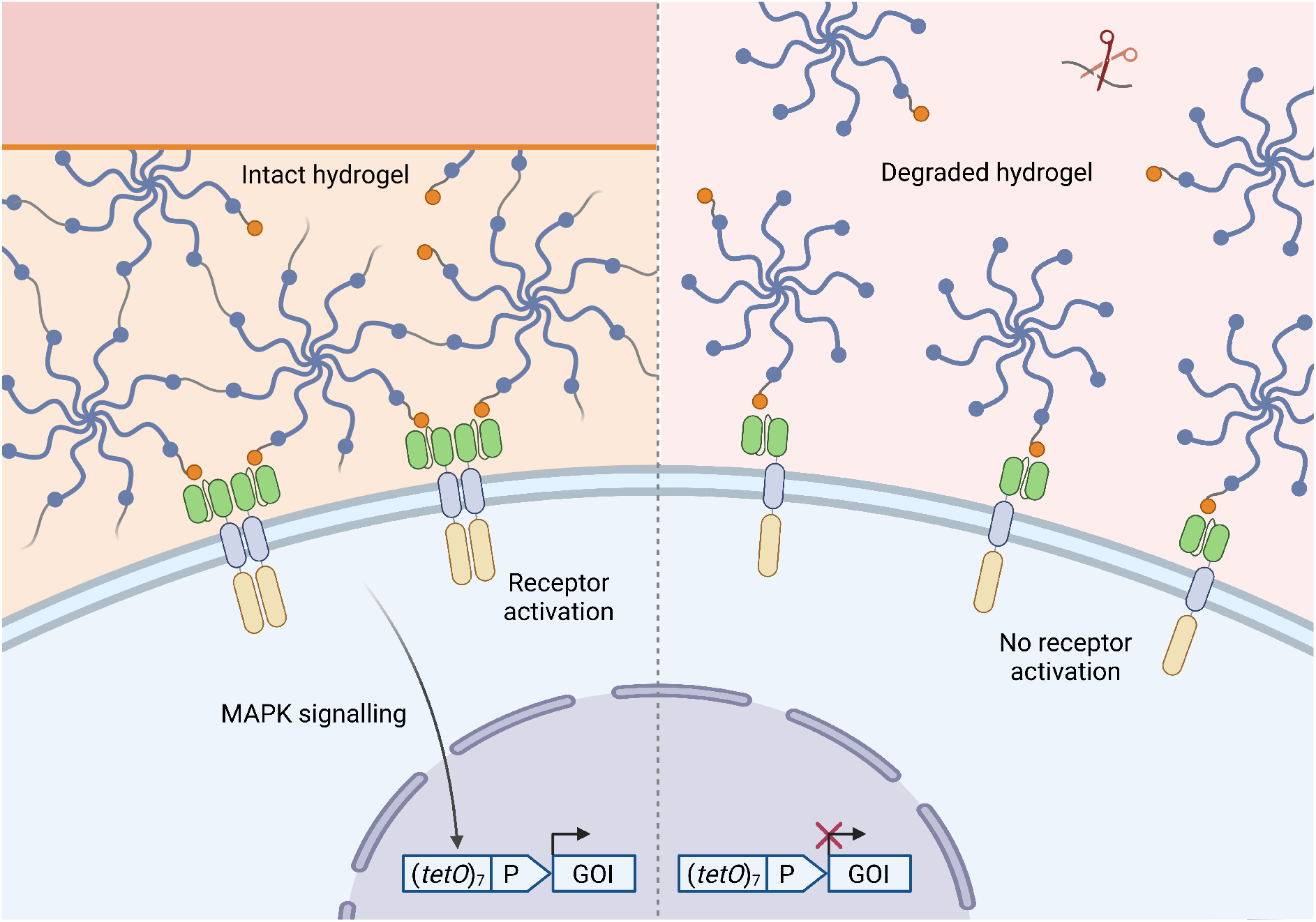
Design of the material-genetic interface to link the presence of an intact, embedding hydrogel to the expression of a target gene. Left: an intact, crosslinked gel contains several fluorescein moieties (orange circles) in close proximity. Upon binding to the synthetic receptors on the cell surface, clustered fluorescein causes the dimerization of the receptors, which in turn triggers MAPK signaling and expression of the gene of interest (GOI). Right: 3C protease (3CP) addition results in the breaking of the crosslinks between PEG units. Therefore, the hydrogel dissolves and the individual fluorescein moieties are unable to cause receptor dimerization, and thus downstream signaling and GOI expression do not take place. *tetO*, tetracycline-responsive operator; P, minimal human cytomegalovirus promoter.

**Figure 2.**
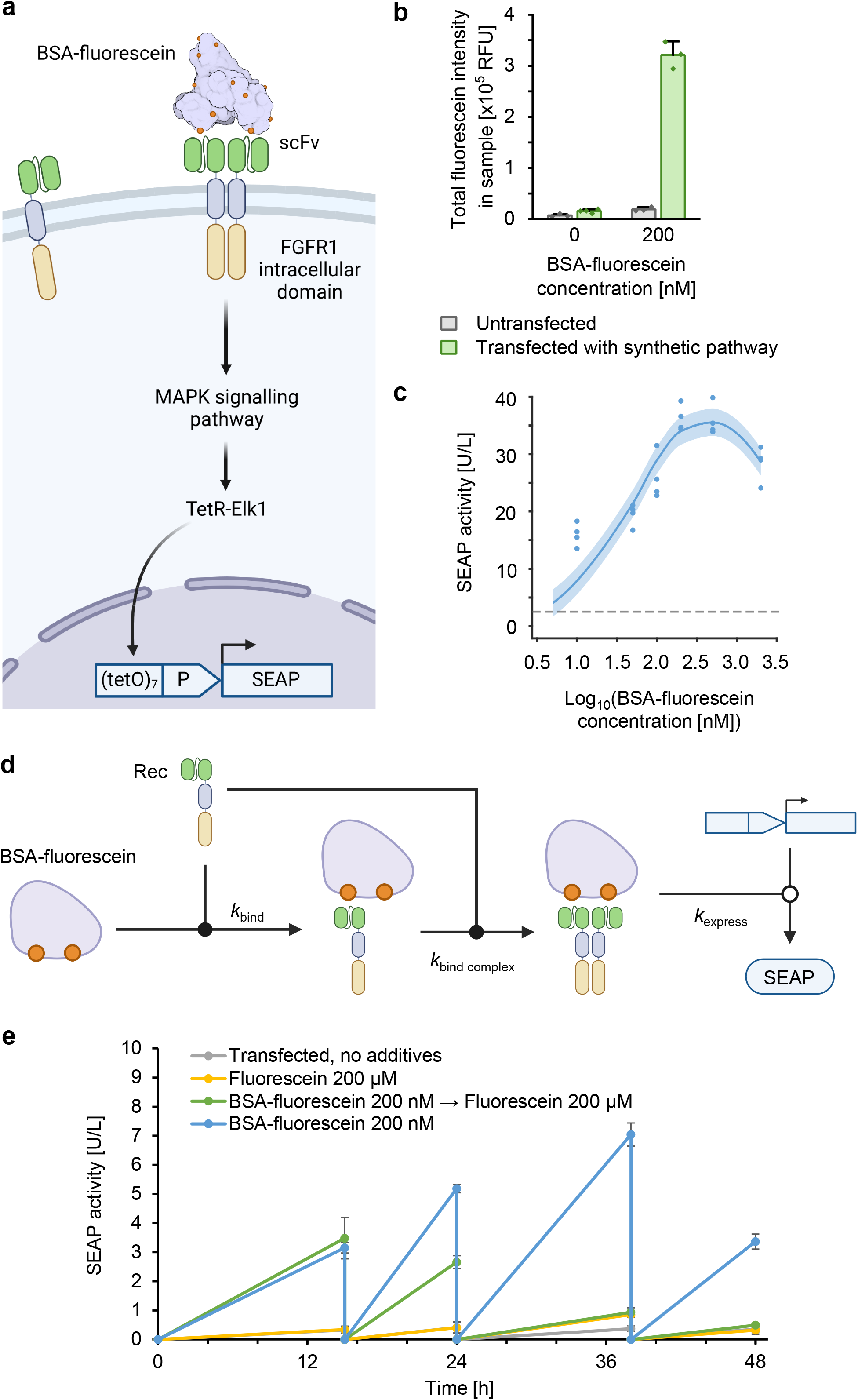
Functional characterization of the fluorescein-responsive signaling pathway. **(a)** Experimental setup. The fluorescein-responsive signaling pathway is activated by BSA-bound clustered fluorescein, which results in SEAP reporter gene expression. SEAP is secreted to the cell culture medium, where its activity can be quantified. **(b)** Evaluation of the production of the fluorescein-responsive receptor at the cell surface. HEK-293T cells were transfected with the expression vector for the fluorescein-responsive receptor (pCJL503), TetR-Elk1 (MKp37) and the SEAP reporter (pMF111), and incubated with 200 nM BSA-fluorescein for 16 h prior to analysis by flow cytometry. The integrated fluorescein intensity per sample is shown. **(c)** Dose-response behavior of the fluorescein-responsive signaling pathway. HEK-293T cells were transfected with expression vectors for the fluorescein-responsive receptor (pCJL503), TetR-Elk1 (MKp37) and the SEAP reporter (pMF111). Cells were subsequently incubated for 5 h prior to addition of the indicated BSA-fluorescein concentrations, cultivation for another 20 h and measurement of SEAP activity. The blue curve represents the model fit to the data. The error band indicates the experimental error of the measurements. The dotted grey line indicates the SEAP activity value in the absence of BSA-fluorescein, which was used as an offset for the model. A detailed description of the model can be found in the Supplementary Material. **(d)** Model graph for the dynamical ordinary differential equation model. BSA-fluorescein binds to a single fluroescein-responsive receptor (Rec) with the rate *k*_bind_. This complex then binds to another receptor with the rate *k*_bind complex_. The BSA-bound receptor dimer then induces expression of SEAP with the rate *k*_express_. **(e)** Characterization of the reversibility of the fluorescein-responsive signaling pathway. HEK-293T cells transfected with the pathway components (**Fig. 1c**) were incubated with 200 nM BSA-fluorescein for 15 h, after which the cell culture medium was exchanged and 200 μM free fluorescein or 200 nM BSA-fluorescein was added. SEAP production was quantified at several timepoints over another 33 h, replacing the cell culture medium with fresh DMEM complete medium with BSA-fluorescein or free fluorescein after each measurement. As negative controls, transfected cells were incubated without addition of any compounds or in the presence of free fluorescein (200 μM). The results shown here correspond to three replicates per sample.

**Figure 3.**
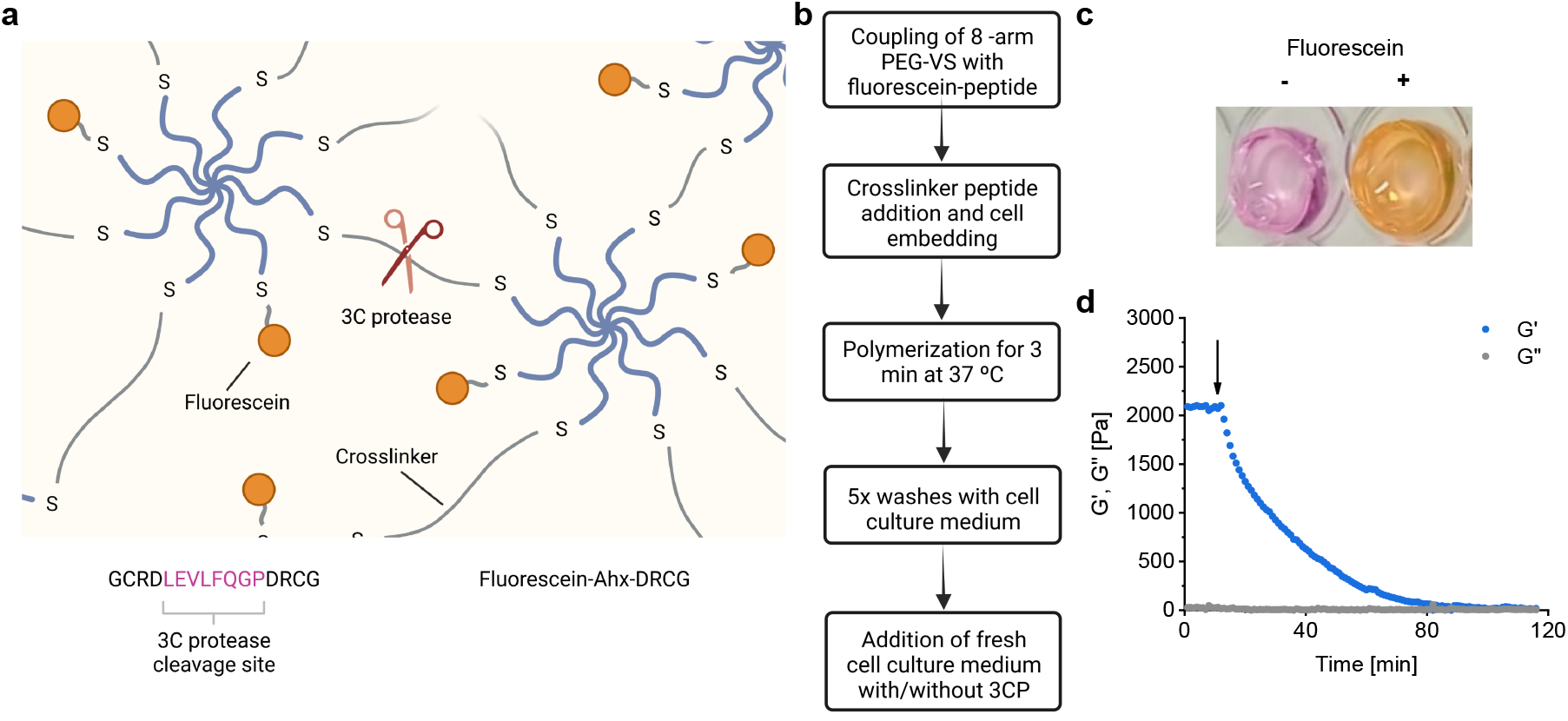
Synthesis and characterization of the hydrogels for cell embedding. **(a)** Hydrogel structure and composition. 8-arm PEG-VS was crosslinked to form a hydrogel using peptides containing two cysteines via Michael-type addition. The peptide further contained a cleavage site for 3CP between the cysteine residues to allow degradation of the hydrogel with 3CP. To functionalize the hydrogel with clustered fluorescein, PEG-VS was coupled to sub-stochiometric amounts of a cysteine-containing peptide conjugated to fluorescein. Bottom left: crosslinker peptide. Bottom right: fluorescein-conjugated peptide. Ahx, aminohexanoic acid. **(b)** Workflow for hydrogel synthesis and embedding of cells. **(c)** Hydrogels with and without the fluorescein-conjugated peptide, retrieved from the cell culture wells after incubation over night in cell-culture medium. **(d)** Rheological characterization of PEG-fluorescein hydrogels. Hydrogels were synthesized containing a fluorescein:VS ratio of 1:10 and subjected to small amplitude oscillatory shear rheology to determine the storage (G’) and loss (G’’) modulus. After 10 min (arrow), the gel was surrounded by a solution containing 0.3 mg/mL 3CP. The shown data are representative for 3 independent experiments.

**Figure 4.**
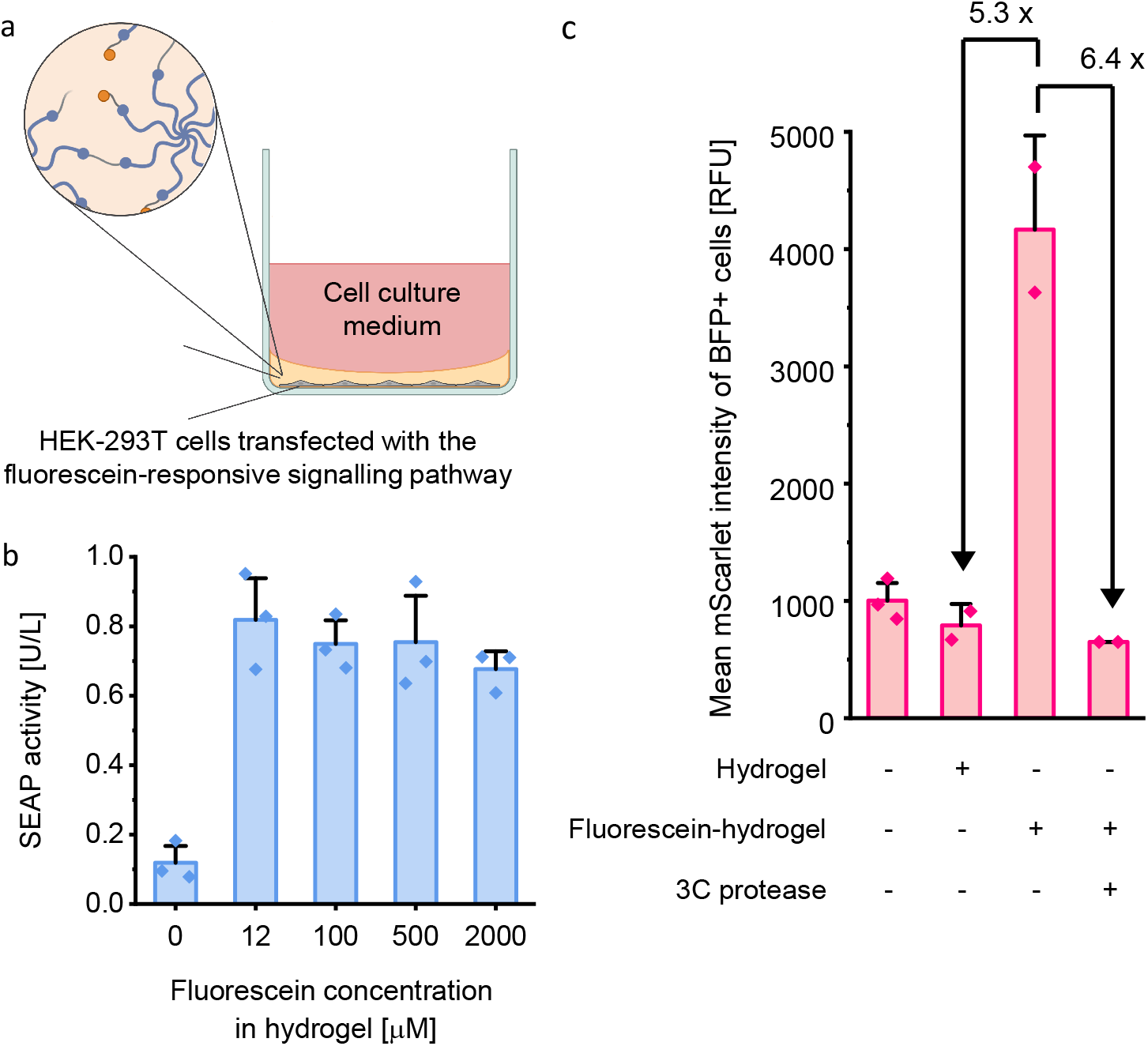
Functional characterization of the material-genetic interface. **(a)** Experimental set-up. HEK-293T cells transfected with the fluorescein-responsive signaling system were embedded in the hydrogel displaying clustered fluorescein. **(b)** HEK-293T cells were transfected with the expression vectors for the fluorescein-responsive chimeric receptor (pCJL503), TetR-Elk1 (MKp37) and the tetracycline-responsive SEAP expression vector (pMF111). Cells were embedded in PEG-based hydrogels functionalized with the indicated concentrations of fluorescein. After cultivation for 20 h, SEAP production was quantified in the supernatant. **(c)** HEK-293T cells were transfected with the expression vectors for the fluorescein-responsive chimeric receptor (pCJL503), TetR-Elk1 (MKp37), the tetracycline-responsive mScarlet expression vector (pCJL510) and the constitutive BFP expression vector (pOK045). Cells were embedded in PEG-based hydrogels containing 500 μM fluorescein. The embedded cells were cultivated in the absence or presence (0.3 mg/mL) of 3CP for 20 h prior to quantifying mScarlet production by flow cytometry. As controls, transfected cells were incubated either in the absence of a hydrogel or embedded in a hydrogel without fluorescein functionalization. The mean mScarlet fluorescence intensity of the BFP-positive cells (as marker for transfected cells) is shown. The results shown here are representative of three independent experiments.

## 3 Results and discussion

### 3.1 Design of the material-genetic interface

We designed a material-genetic interface to functionally link the presence of an intact, embedding hydrogel to the expression of a desired transgene. This interface consists of two components, first, clustered ligands in the embedding hydrogel and second, a synthetic cell surface receptor that is activated by these ligands and subsequently triggers a signaling cascade to induce expression of a target gene (**Fig. 1**). Upon hydrogel degradation, the clustering of the ligands is reverted thus resulting in de-activation of the receptor and downstream transgene expression. Similarly, in cells growing outside the hydrogel, the receptor is de-activated. We selected fluorescein as a ligand, which is clinically licensed as an intravenously applied contrast agent and shows an excellent safety profile^43^. Fluorescein can easily be conjugated to hydrogel components or other biomolecules containing primary amines using FITC. To genetically sense clustered fluorescein, we engineered a synthetic receptor based on the GEMS platform^38^. To this aim, we fused an scFv specific for fluorescein (FITC-E2^39^) via the erythropoietin receptor (EpoR) transmembrane domain to the intracellular domain of the fibroblast growth factor receptor 1 (FGFR1). Ligand-induced clustering of the receptor results in FGFR signaling, the activation of the mitogen-activated protein kinase (MAPK) pathway and the nuclear translocation of a synthetic transcription factor composed of ETS-like protein 1 (Elk1) fused to the bacterial tetracycline repressor TetR (TetR-Elk1). Nuclear TetR-Elk1 binds to the tetracycline-responsive operator *(tetO)* and induces the adjacent minimal human cytomegalovirus promoter P_hCMVmin_, resulting in expression of the downstream target gene^38^.

### 3.2 Functional characterization of the synthetic fluorescein-responsive signaling pathway

To evaluate whether the fluorescein-specific scFv is expressed at the cell surface and able to bind fluorescein, we transfected HEK-293T cells with the fluorescein-responsive synthetic receptor (pCJL503) together with a TetR-Elk1 expression cassette (MKp37) and a tetracycline-responsive expression plasmid (pMF111) for the human placental SEAP reporter, and incubated the cells with fluorescein-conjugated BSA prior to flow cytometry analysis (**Fig. 2a**). We analyzed the average fluorescein fluorescence per transfected cell and found it to be up to 46 times higher in transfected cells incubated with BSA-fluorescein as compared to non-transfected cells or cells incubated with buffer only, confirming the presence of the synthetic receptors on the plasma membrane and the functionality of the scFv (**Fig. 2b**, see **Fig. S1** for flow cytometry gating strategy and **Table S3** for raw data). To evaluate functionality of the complete system, we transfected HEK-293T cells with the fluorescein-responsive pathway as described above. To stimulate the receptor via clustered fluorescein, we added fluorescein-conjugated BSA at concentrations of 10 – 2000 nM (**Fig. 2c**). By measuring SEAP production after 20 h, we found a dose-dependent increase in SEAP levels for BSA-fluorescein concentrations of 10 – 200 nM indicating the functionality of the synthetic receptor and downstream signaling cascade. At higher fluorescein concentrations, slightly decreased SEAP values were observed. We hypothesize that at high BSA-fluorescein concentrations the ligands compete for receptor binding. Thus, only one BSA-fluorescein might be bound per receptor which would not result in receptor crosslinking and downstream signaling. To support this hypothesis, we have developed a dynamical mathematical model based on ordinary differential equations that reflects the competition of the ligands for the receptor (**Fig. 2d**, for the model description, see Supplementary Material (**Figures S5**, **S6** and **S7** and **Table S5**)). The model was shown to explain the observed shape of the dose-response curve (**Fig. 2c**).

Finally, we tested the reversibility of the fluorescein-responsive signaling pathway, which would be required for gene expression to turn off when the cells lose contact with the hydrogel. For this purpose, we incubated transfected HEK-293T cells with 200 nM BSA-fluorescein for 15 h. We subsequently replaced the medium by fresh medium either containing free fluorescein (200 μM) or BSA-fluorescein (200 nM), and monitored SEAP activity at several intervals over another 33 h. The cell culture medium was replaced with fresh medium with BSA-fluorescein or free fluorescein after each measurement. (**Fig. 2e**). In samples containing free fluorescein after an initial incubation step with BSA-fluorescein, SEAP production decreased over time back to the level of the samples incubated only with fluorescein, whereas samples containing BSA-fluorescein throughout the whole experiment showed increased SEAP titers after every incubation period. In contrast, cells cultivated without fluorescein or with free fluorescein for the whole experiment showed background SEAP levels only. These findings indicate that the activation of the receptor is strictly dependent on clustered fluorescein and that the activation is reversible, both of which are important for the functionality of the material-genetic interface as a safety switch for encapsulated cell therapy.

### 3.3 Synthesis and characterization of hydrogels displaying clustered fluorescein for cell encapsulation

As model material for cell encapsulation, we used a PEG-based hydrogel due to the high cell compatibility of this polymer and its broad usage in cell encapsulation^44^. As precursor for the hydrogel, we used 8-arm PEG (40 kDa) terminally functionalized with vinylsulfone (VS) groups. For hydrogel formation, the VS groups were reacted via Michael-type addition with the thiol groups of peptides harboring two cysteines. To emulate degradation of the embedding hydrogel, the peptides further harbored a cleavage site for the 3CP between the two cysteines (**Fig. 3a**). To incorporate clustered fluorescein into the hydrogel, PEG-VS was reacted with fluorescein-labeled peptides containing a cysteine residue prior to crosslinking. The hydrogel can directly be polymerized in the presence of cells (**Fig. 3b-c, Fig. S4**), similarly to previous studies embedding cells into PEG-VS-based hydrogels^45^. We characterized the resulting gels by small amplitude oscillatory shear rheology revealing a storage modulus (G’) of 2.1 kPa (**Fig. 3d**) which is in the range of previous hydrogels used for cell encapsulation^45^. We finally evaluated whether the hydrogels can be degraded by the addition of 3CP. To this aim, the gel was surrounded by a solution containing 3CP (10 min after starting the measurement, see arrow in **Fig. 3d**). We subsequently observed a strong decrease in G’ down to the level of the loss modulus (G’’) after ~90 min, indicating the dissolution of the hydrogel.

### 3.4 Assembly and functional validation of the material-genetic interface

In order to functionally validate the material-genetic interface, we combined cells engineered with the fluorescein-responsive signaling pathway with the hydrogel material displaying clustered fluorescein, and validated target gene expression in the presence of an intact or degraded hydrogel. To this aim, HEK-293T cells were transfected with expression vectors for the fluorescein-responsive chimeric receptor (pCJL503), TetR-Elk1 (MKp37) and the tetracycline-responsive SEAP expression vector (pMF111). Cells were subsequently embedded in PEG-based hydrogels functionalized with different ratios of fluorescein-labeled peptide:VS ranging from 1:833 to 1:5 corresponding to fluorescein concentrations of 12 μM to 2 mM in the final hydrogel, respectively (**Fig. 4a**). The cells were cultivated for 20 h in the hydrogel prior to analysis of SEAP production. We observed that there were no significant differences in reporter gene expression throughout the tested fluorescein concentrations in the hydrogel (**Fig. 4b**). For further experiments, we prepared hydrogels with a fluorescein concentration of 500 μM.

Finally, we investigated whether degradation of the embedding hydrogel would effectively de-activate transgene expression. In these experiments we replaced the SEAP reporter by the fluorescent reporter mScarlet (pCJL510) in order to avoid possible effects of hydrogel integrity (intact/degraded) on diffusion and release of the secreted SEAP reporter, as well as effects of hydrogel or protease incubation on cell growth. For normalization, we co-transfected an expression vector for constitutive BFP (pOK045). Cells were subsequently embedded into PEG-based hydrogels functionalized with 500 μM fluorescein. The samples were cultivated in the presence or absence of 3CP for 20 h prior to quantifying mScarlet and BFP production (**Fig. 4c,** see **Fig. S2** for flow cytometry gating strategy and **Table S4** for raw data). In hydrogel-embedded cells we observed high mScarlet production in accordance with the data obtained using the SEAP reporter (**Fig. 4b**). However, upon addition of 3CP and subsequent hydrogel degradation, mScarlet levels returned to baseline levels, as observed in cells embedded in hydrogels without fluorescein functionalization or in cells cultivated in the absence of an embedding hydrogel (**Fig. 4c**). These data demonstrate that the material-genetic interface functionally links the presence of an embedding hydrogel to the expression of the target gene.

## 4 Conclusion and outlook

In the present study, we have engineered mammalian cells with a synthetic pathway, based on the GEMS system, in order to give them the ability to sense a safe and biocompatible moiety present in an embedding hydrogel, but not occurring naturally in the human body. The cells are able to sense whether or not they are in contact with the hydrogel, and tune target gene expression accordingly. The strategy presented here can in principle be broadly applied to any cell type, material or extracellular ligand.

We envision that this tool could be applied in encapsulated cell therapy in two different ways. In a combined therapeutic gene expression control and safety switch approach, the gene of interest could be replaced by a gene that results in the production of the therapeutic agent. As a result, if the engineered cells escape from the encapsulation material, they will cease to produce this agent. This strategy would be useful for therapies in which off-target toxicities are a concern, and the production of the drug should be confined to its intended site of action. On the other hand, the genetic network controlling the safety switch could be used orthogonally to another genetic network controlling the production of the therapeutic agent. One way of doing this would be by using cells lacking an essential synthesis pathway. In this case, the reporter would be replaced by a gene coding for an enzyme that compensates for the missing pathway. Consequently, cells escaping the hydrogel would not receive the necessary stimulus to transcribe this survival gene, and would not survive outside the hydrogel. This approach could prove useful for therapies in which the presence of the therapeutic cells outside the encapsulation material constitutes a safety concern, for example due to immunogenicity.

Finally, so as to apply this design to a wide range of encapsulated cell therapies, a stable insertion of DNA would be required instead of transient transfection. For such purposes, different systems have been described that allow the stable installation of multiple genes in different cell lines^46–48^. In addition, since the final product would involve genetically modified cells, it would have to fulfill the biosafety requirements that apply to genetically modified organisms used in the clinic^49^.

This constitutes the first step towards building a novel safety switch for encapsulated therapeutic cells that addresses the issue of escape of the engineered cells from the encapsulation material due to proliferation or material damage. The switch could contribute to prevent side effects (such as off-target toxicities or host immune system activation), without the need for a medical intervention or the termination of the therapy, and without irreversibly killing all therapeutic cells, allowing the treatment to proceed for longer.

## 5 Acknowledgements

We would like to thank Prof. Martin Fussenegger, ETH Zurich for kindly providing plasmids pLeo669 and MKp37. We are grateful to Oliver Thomas, Katharina Ostmann and Hanna Wagner for the pOT21, pKO045 and pHJW257 plasmids, respectively, as well as to Nadine Urban for the manufacturing of the ring for rheology measurements. We also thank Hanyang Cai for her contribution to the project during her internship. This work was supported by the German Research Foundation (Deutsche Forschungsgemeinschaft, DFG) under Germany’s Excellence Strategy – CIBSS, EXC-2189, Project ID: 390939984 and under the Excellence Initiative of the German Federal and State Governments – BIOSS, EXC-294 and SGBM, GSC-4, and in part by the Ministry for Science, Research and Arts of the State of Baden-Württemberg.

## Notes

https://github.com/fgwieland/Material-genetic-interface-model

### Competing Interest Statement

The authors have declared no competing interest.

