## Supplementaru for "Engineering a material-genetic interface as safety switch for embedded therapeutic cells"

### SUPPLEMENTARY MATERIAL

**Table S1. Plasmids generated in this study**

| Name | Description | Sequence |
|---|---|---|
| **pCJL503** | SV40 promoter-IgK secretion signal-scFv(Flu-E2)-EpoR-FGFR1<br><br>Backbone: pcDNA3.1 | ...ATATACGCGTGATCTGCGATCTGCATCTCAATTAGTCAGCAACCATAGTCCC<br>GCCCCTAACTCCGCCCATCCCGCCCCTAACTCCGCCCAGTTCCGCCCATTCTCC<br>GCCCCATCGCTGACTAATTTTTTTTATTTATGCAGAGGCCGAGGCCGCCTCGG<br>CCTCTGAGCTATTCCAGAAGTAGTGAGGAGGCTTTTTTGGAGGCCTAGGCTTT<br>TGCAAAAAGCTTCGAATCGCGCTAGCGGCCGGCCACCATGGAAACTGATACT<br>TTGCTGCTCTGGGTCCTGCTGCTGTGGGTCCCTGGATCAACGGGGGACCAGG<br>TGCAGCTGGTGGAGTCTGGGGGAAACTTGGTACAGCCTGGGGGGTCCCTGA<br>GACTCTCCTGTGCAGCCTCTGGATTCACCTTTGGCAGCTTTGCCATGAGCTGG<br>GTCCGCCAGGCTCCAGGGGGGGGGCTGGAGTGGGTCGCAGGTCTGAGTGCT<br>CGTAGTAGTCTCACACACTATGCAGACTCCGTGAAGGGCCGGTTCACCATCTC<br>CAGAGACAACGCCAAGAACTCAGTGTATCTGCAAATGAACAGCCTGAGAGTC<br>GAGGACACGGCTGTGTATTACTGTGCGAGAAGATCGTATGATAGTAGTGGTT<br>ATTGGGGCCACTTCTACTCCTACATGGACGTCTGGGGCCAAGGCACCCTGGTC<br>ACCGTCTCGAGTGGTGGAGGCGGTTCAGGCGGAGGTGGCAGCGGCGGTGGC<br>GGATCGCAGTCTGTGTTGACGCAGCCGTCCTCAGTGTCTGCGGCCCCAGGAC<br>AGAAGGTCACCATTTCCTGCTCTGGAAGCACCTCCAACATTGGGAATAATTAT<br>GTCTCCTGGTACCAACAGCACCCAGGCAAAGCCCCCAAACTCATGATTTATGA<br>TGTCAGTAAGCGGCCCTCAGGGGTCCCTGACCGATTCTCTGGCTCCAAGTCTG<br>GCAACTCAGCCTCCCTGGACATCAGTGGGCTCCAGTCTGAGGATGAGGCTGA<br>TTATTACTGTGCAGCATGGGATGACAGCCTGAGTGAATTTCTCTTCGGAACTG<br>GGACCAAGCTGACCGTCCTAGGGGCCTCGGGGGCCGATAGCTCCGGAGAATT<br>CGCACCTTCACCCAGCCTCCCGGACCCCAAGTTTGAGAGCAAAGCGGCCCTGC<br>TGGCATCCCGGGGCTCCGAAGAACTTCTGTGCTTCACCCAACGCTTGGAAGAC<br>TTGGTGTGTTTCTGGGAGGAAGCGGCGAGCTCCGGGATGGACTTCAACTACA<br>GCTTCTCATACCAGCTCGAGGGTGAGTCACGAAAGTCATGTAGCCTGCACCAG<br>GCTCCCACCGTCCGCGGCTCCGTGCGTTTCTGGTGTTCACTGCCAACAGCGGA<br>CACATCGAGTGCTGTGCCGCTGGAGCTGCAGGTGACGGAGGCGTCCGGTTCT<br>CCTCGCTATCACCGCATCATCCATATCAATGAAGTAGTGCTCCTGGACGCCCCC<br>GCGGGGCTGCTGGCGCGCCGGGCAGAAGAGGGCAGCCACGTGGTGCTGCGC<br>TGGCTGCCACCTCCTGGAGCACCTATGACCACCCACATCCGATATGAAGTGGA<br>CGTGTCGGCAGGCAACCGGGCAGGAGGGACACAAAGGGTGGAGGTCCTGG<br>AAGGCCGCACTGAGTGTGTTCTGAGCAACCTGCGGGGCGGGACGCGCTACAC<br>CTTCGCTGTTCGAGCGCGCATGGCCGAGCCGAGCTTCAGCGGATTCTGGAGT<br>GCCTGGTCTGAGCCCGCGTCACTACTGACCGCTAGCGACCTGGACCCTCTCAT<br>CTTGACGCTGTCTCTCATTCTGGTCCTCATCTCGTTGTTGCTGACGGTTCTGGC<br>CCTGCTGTCCATGAAGAGCGGCACCAAGAAGAGCGACTTCCATAGCCAGATG<br>GCTGTGCACAAGCTGGCCAAGAGCATCCCTCTGCGCAGACAGGTAACAGTGT<br>CAGCTGACTCCAGTGCATCCATGAACTCTGGGGTTCTCCTGGTTCGGCCCTCA<br>CGGCTCTCCTCCAGCGGGACCCCCATGCTGGCTGGAGTCTCCGAATATGAGCT<br>CCCTGAGGATCCCCGCTGGGAGCTGCCACGAGACAGACTGGTCTTAGGCAAA<br>CCACTTGGCGAGGGCTGCTTCGGGCAGGTGGTGTTGGCTGAGGCCATCGGGC<br>TGGATAAGGACAAACCCAACCGTGTGACCAAAGTGGCCGTGAAGATGTTGAA<br>GTCCGACGCAACGGAGAAGGACCTGTCGGATCTGATCTCGGAGATGGAGAT<br>GATGAAAATGATTGGGAAGCACAAGAATATCATCAACCTTCTGGGAGCGTGC<br>ACACAGGATGGTCCTCTTTATGTCATTGTGGAGTACGCCTCCAAAGGCAATCT<br>CCGGGAGTATCTACAGGCCCGGAGGCCTCCTGGGCTGGAGTACTGCTATAAC |

| | | |
|---|---|---|
| | | CCCAGCCACAACCCCGAGGAACAGCTGTCTTCCAAAGATCTGGTATCCTGTGC<br>CTATCAGGTGGCTCGGGGCATGGAGTATCTTGCCTCTAAGAAGTGTATACACC<br>GAGACCTGGCTGCTAGGAACGTCCTGGTGACCGAGGATAACGTAATGAAGAT<br>CGCAGACTTTGGCTTAGCTCGAGACATTCATCATATCGACTACTACAAGAAAA<br>CCACCAACGGCCGGCTGCCTGTGAAGTGGATGGCCCCTGAGGCGTTGTTTGA<br>CCGGATCTACACACACCAGAGCGATGTGTGGTCTTTTGGAGTGCTCTTGTGGG<br>AGATCTTCACTCTGGGTGGCTCCCCATACCCCGGTGTGCCTGTGGAGGAACTT<br>TTCAAGCTGCTGAAGGAGGGTCATCGAATGGACAAGCCCAGTAACTGTACCA<br>ATGAGCTGTACATGATGATGCGGGACTGCTGGCATGCAGTGCCCTCTCAGAG<br>ACCTACGTTCAAGCAGTTGGTGGAAGACCTGGACCGCATTGTGGCCTTGACCT<br>CCAACCAGGAGTATCTGGACCTGTCCATACCGCTGGACCAGTACTCACCCAGC<br>TTTCCCGACACACGGAGCTCCACCTGCTCCTCAGGGGAGGACTCTGTCTTCTCT<br>CATGAGCCGTTACCTGAGGAGCCCTGTCTGCCTCGACACCCCACCCAGCTTGC<br>CAACAGTGGACTCAAACGGCGCTAGGCGGCCGCTC… |
| **pCJL510** | Tetracycline-inducible minimal CMV promoter-mScarlet<br><br>Backbone: pTBC1 | …CCTTTCGTCCTCGAGTTTACCACTCCCTATCAGTGATAGAGAAAAGTGAAAG<br>TCGAGTTTACCACTCCCTATCAGTGATAGAGAAAAGTGAAAGTCGAGTTTACC<br>ACTCCCTATCAGTGATAGAGAAAAGTGAAAGTCGAGTTTACCACTCCCTATCA<br>GTGATAGAGAAAAGTGAAAGTCGAGTTTACCACTCCCTATCAGTGATAGAGA<br>AAAGTGAAAGTCGAGTTTACCACTCCCTATCAGTGATAGAGAAAAGTGAAAG<br>TCGAGTTTACCACTCCCTATCAGTGATAGAGAAAAGTGAAAGTCGAGCTCGGT<br>ACCCGGGTCGAGTAGGCGTGTACGGTGGGAGGCCTATATAAGCAGAGCTCGT<br>TTAGTGAACCGTCAGATCGCCTGGAGACGCCATCCACGCTGTTTTGACCTCCA<br>TAGAAGACACCGGGACCGATCCAGCCTCCGCGGCCCCGAATTCGAGCTCGCC<br>CGGGGATCCTCTAGAGTCAGCTTCTGCATGGTGAGCAAGGGCGAGGCAGTGA<br>TCAAGGAGTTCATGCGGTTCAAGGTGCACATGGAGGGCTCCATGAACGGCCA<br>CGAGTTCGAGATCGAGGGCGAGGGCGAGGGCCGCCCCTACGAGGGCACCCA<br>GACCGCCAAGCTGAAGGTGACCAAGGGTGGCCCCCTGCCCTTCTCCTGGGAC<br>ATCCTGTCCCCTCAGTTCATGTACGGCTCCAGGGCCTTCACCAAGCACCCCGCC<br>GACATCCCCGACTACTATAAGCAGTCCTTCCCCGAGGGCTTCAAGTGGGAGCG<br>CGTGATGAACTTCGAGGACGGCGGCGCCGTGACCGTGACCCAGGACACCTCC<br>CTGGAGGACGGCACCCTGATCTACAAGGTGAAGCTCCGCGGCACCAACTTCC<br>CTCCTGACGGCCCCGTAATGCAGAAGAAGACAATGGGCTGGGAAGCGTCCAC<br>CGAGCGGTTGTACCCCGAGGACGGCGTGCTGAAGGGCGACATTAAGATGGC<br>CCTGCGCCTGAAGGACGGCGGCCGCTACCTGGCGGACTTCAAGACCACCTAC<br>AAGGCCAAGAAGCCCGTGCAGATGCCCGGCGCCTACAACGTCGACCGCAAGT<br>TGGACATCACCTCCCACAACGAGGACTACACCGTGGTGGAACAGTACGAACG<br>CTCCGAGGGCCGCCACTCCACCGGCGGCATGGACGAGCTGTACAAGTAACCC<br>GTGGTCC… |
| **pHJW257** | T7 promoter-StrepTag-3CP<br><br>Backbone: pHJW4[1] | …CCCGCGAAATTAATACGACTCACTATAGGGAGACCACAACGGTTTCCCTCTA<br>GAAATAATTTTGTTTAACTTTAAGAAGGAGATATACATATGTGGAGCCACCCG<br>CAGTTCGAGAAATCGGCGGGTCCGAATACCGAATTTGCACTGAGCCTGCTGC<br>GTAAAAACATTATGACCATTACCACCTCCAAAGGCGAATTTACCGGTCTGGGT<br>ATTCATGATCGTGTTTGTGTTATTCCGACCCATGCACAGCCTGGTGATGACGTT<br>CTGGTTAATGGTCAGAAAATTCGCGTGAAAGATAAATACAAACTGGTGGATC<br>CGGAAAACATTAATCTGGAACTGACCGTTCTGACCCTGGATCGTAATGAAAAA<br>TTTCGTGATATCCGTGGCTTTATCAGCGAAGATCTGGAAGGTGTTGATGCAAC<br>CCTGGTTGTTCATAGCAATAACTTTACCAACACCATTCTGGAAGTTGGTCCGGT<br>TACCATGGCAGGTCTGATTAATCTGAGCAGCACCCCGACCAATCGTATGATTC<br>GTTATGATTATGCAACCAAAACCGGTCAGTGTGGTGGTGTTCTGTGTGCAACC<br>GGTAAAATCTTTGGCATTCATGTTGGTGGCAATGGTCGTCAGGGTTTTAGCGC<br>ACAGCTGAAAAAACAGTATTTCGTCGAAAAACAGTGAAAGCTTGATC… |

**Table S2. Peptides designed for and used in this study**. Highlighted in blue is the 3C protease cut site. Ahx, aminohexanoic acid.

| Short name | Sequence | Modifications |
|---|---|---|
| **Fluroescein-peptide** | DRCG | N-Terminal Modification: FITC-Ahx<br>C-Terminal Modification: Amidation |
| **Crosslinker peptide** | GCRDLEVLFQGPDRCG | N-Terminal Modification: Acetylation<br>C-Terminal Modification: Amidation |

**a**

**Untransfected HEK-293T cells:**

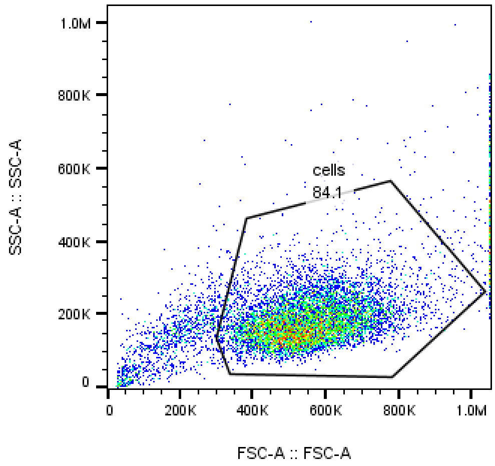


Sample name: Untransfected_1.fcs
Population: Ungated
No. events: 10000

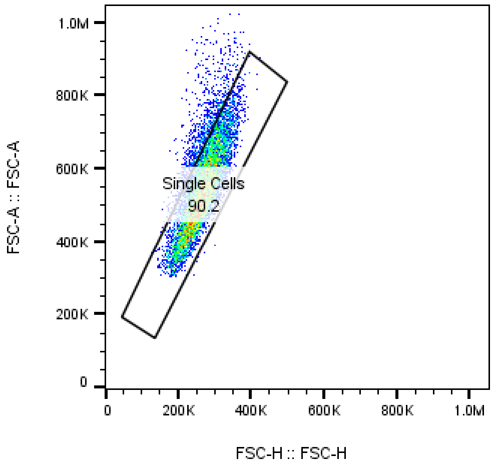


Sample name: Untransfected_1.fcs
Population: cells
No. events: 8407

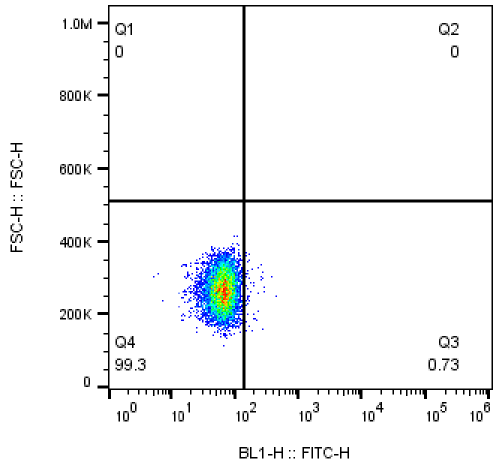


Sample name: Untransfected_1.fcs
Population: Single Cells
No. events: 7582

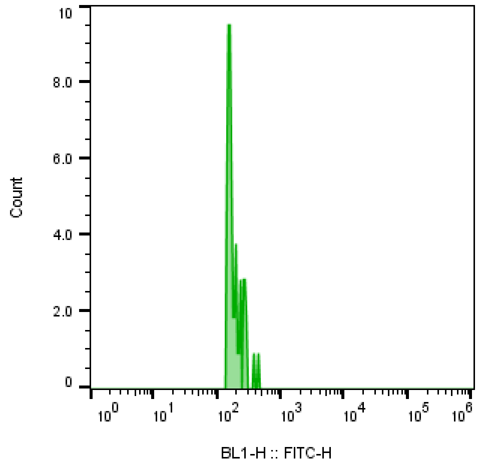


Sample name: Untransfected_1.fcs
Population: Q3: FITC-H+ , FSC-H-
No. events: 55.0

**b**

**HEK-293T cells transfected with the fluorescein-responsive MAPK pathway with 200 nM BSA-fluorescein:**

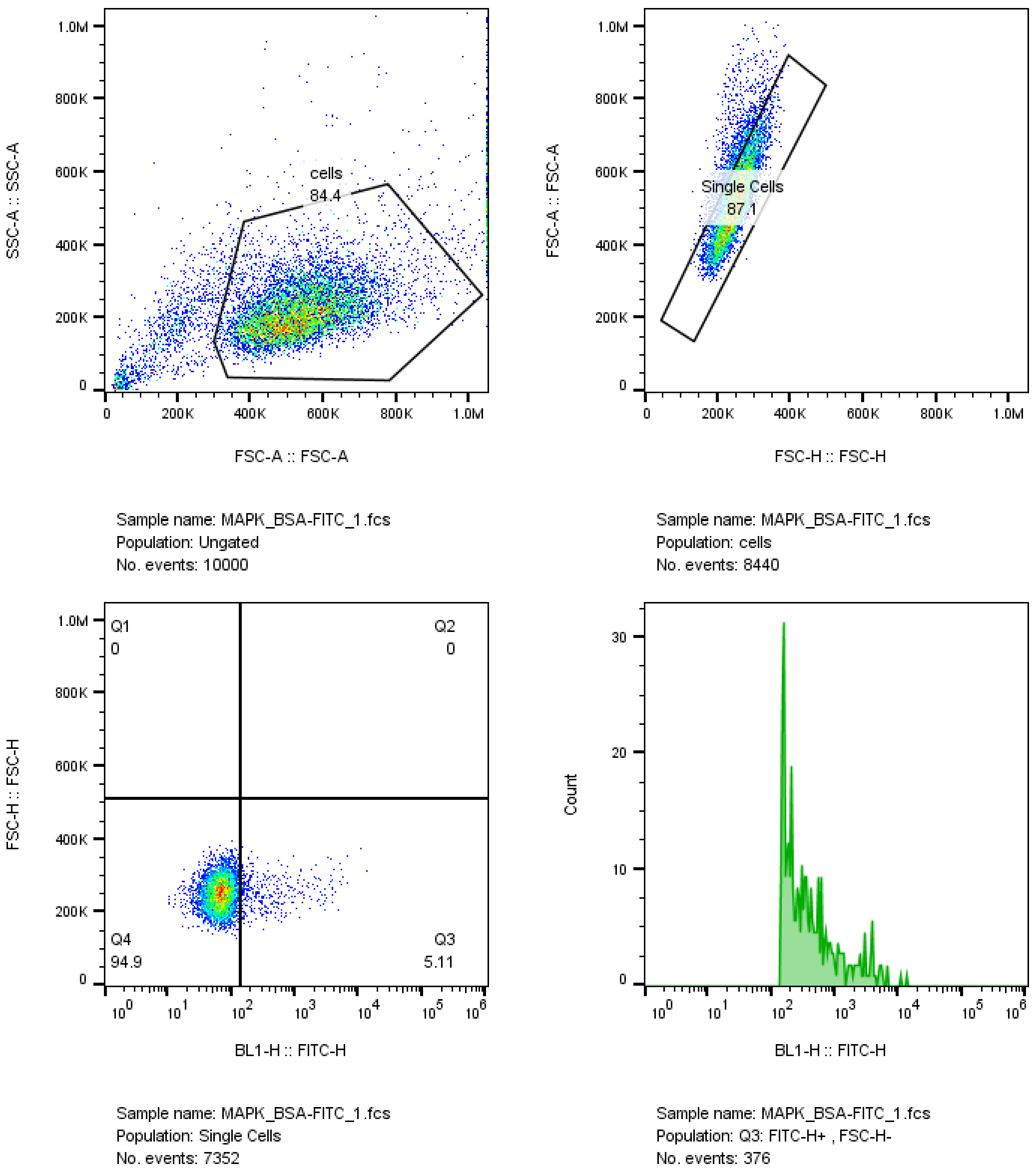


**Figure S1**. **Gating strategy of analysis of flow cytometry data for Fig. 2b**. Fluorescein-positive single cells were gated (shown in histogram) and their intensities were added up. **(a)** Untransfected HEK-293T cells in the absence of BSA-fluorescein. **(b)** HEK-293T cells transfected with the fluorescein-responsive MAPK pathway and incubated with 200 nM BSA-fluorescein. The samples shown here are representative of at least 3 replicates.

**a**

**Untransfected HEK-293T cells, no hydrogel:**

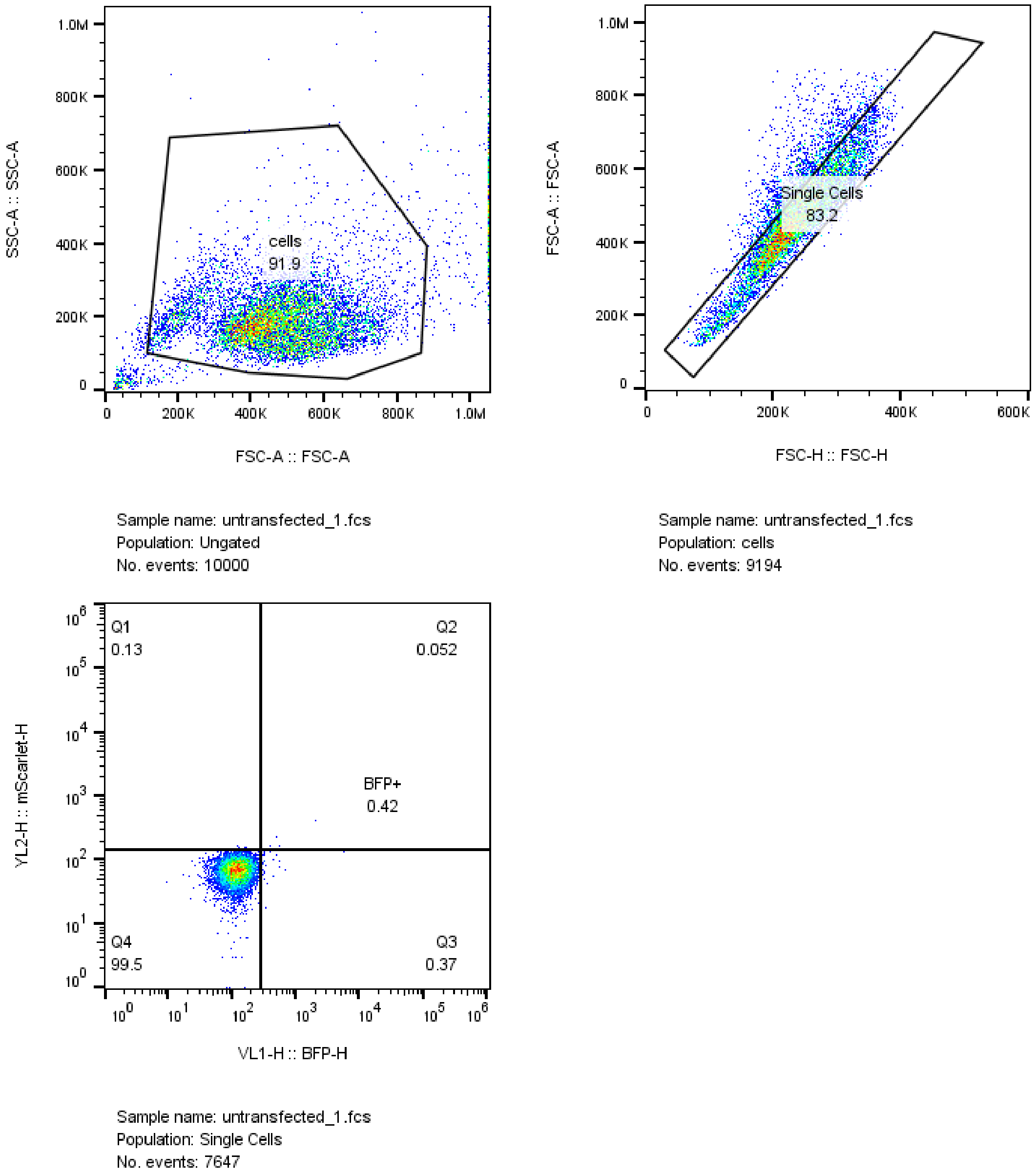

**b**

**HEK-293T cells transfected with the fluorescein-responsive MAPK pathway, embedded in hydrogel, with 3CP:**

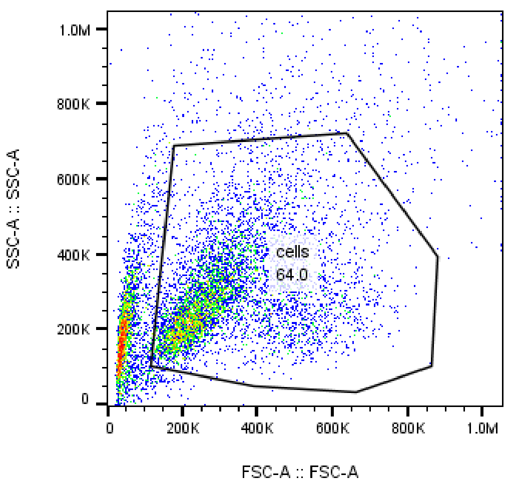


Sample name: with 3CP_1.fcs
Population: Ungated
No. events: 10000

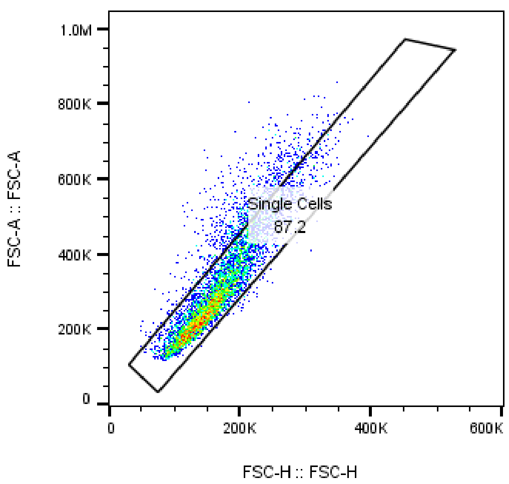


Sample name: with 3CP_1.fcs
Population: cells
No. events: 6403

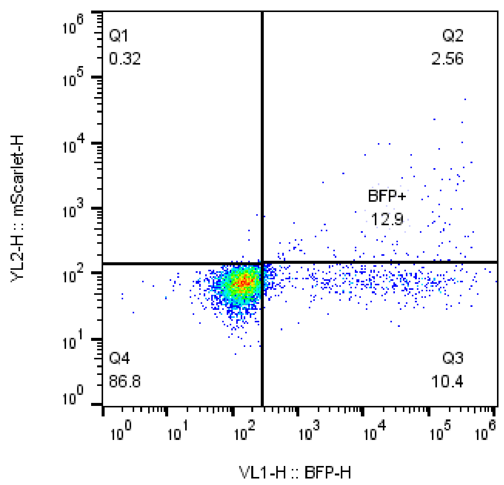


Sample name: with 3CP_1.fcs
Population: Single Cells
No. events: 5582

**c**

**HEK-293T cells transfected with the fluorescein-responsive MAPK pathway, embedded in hydrogel, without 3CP:**

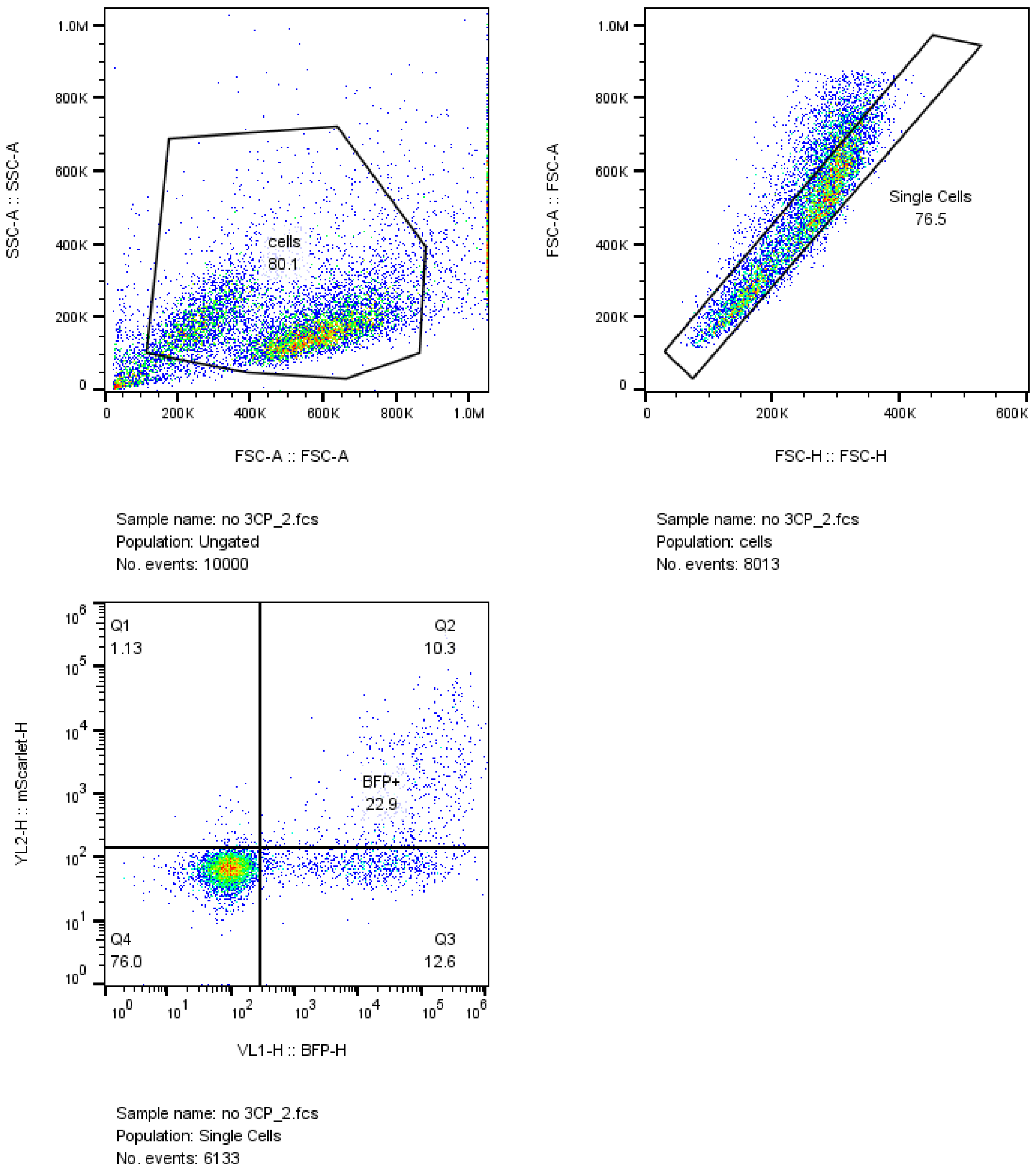


**Figure S2**. **Gating strategy of analysis of flow cytometry data for Fig. 4c**. BFP-positive single cells (quadrants Q2 and Q3) were gated and their mean mScarlet fluorescence intensity was calculated. **(a)** Untransfected HEK-293T cells not embedded in a hydrogel. **(b)** HEK-293T cells transfected with the fluorescein-responsive MAPK pathway, embedded in a hydrogel and incubated with 3CP. **(c)** HEK-293T cells transfected with the fluorescein-responsive MAPK pathway, embedded in a hydrogel and in the absence of 3CP. The samples shown here are representative of at least 2 replicates.

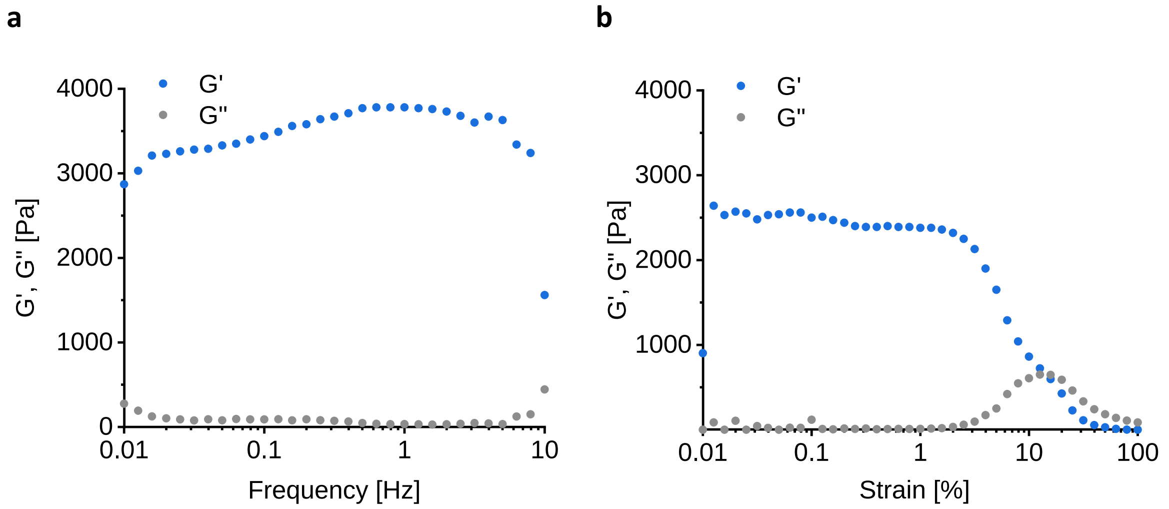


**Figure S3. Mechanical characterization of the hydrogels**. 5% 8-arm PEG-VS was crosslinked to form a hydrogel using peptides containing two cysteines via Michael-type addition. To functionalize the hydrogel with clustered fluorescein, PEG-VS was coupled to a cysteine-containing peptide conjugated to fluorescein, in a 1:10 fluorescein:VS ratio. The storage (G') and loss (G") moduli of the hydrogels were measured by oscillatory shear rheology at frequencies from 0.01 to 10 Hz **(a)** and strain values from 0.01 to 100% **(b)**.

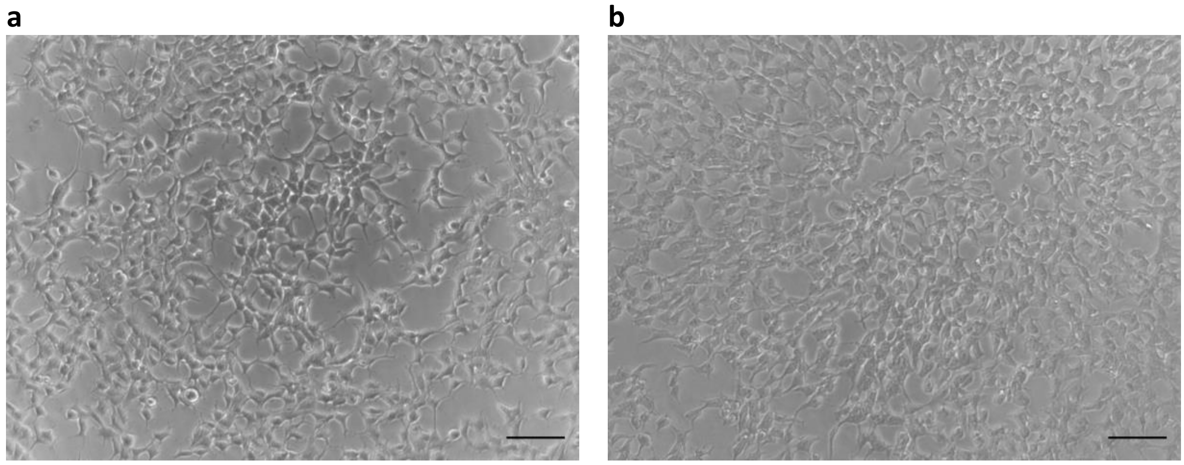


**Figure S4. Microscopy images of cells seeded onto 96-well tissue culture plates (a) and cells seeded onto 96-well tissue culture plates and embedded into fluorescein-containing PEG hydrogels (b).** Scale bar = 100 µm.

**Table S3**

| Sample | Count | cells Count | Single Cells Count | Fluorescein + count | Fluorescein - count | sum of fluroescein int* |
|---|---|---|---|---|---|---|
| MAPK_BSA-FITC_1.fcs | 10000 | 8440 | 7352 | 376 | 6976 | 294013 |
| MAPK_BSA-FITC_3.fcs | 10000 | 8249 | 7232 | 427 | 6805 | 346807 |
| MAPK_BSA-FITC_4.fcs | 10000 | 8574 | 7373 | 407 | 6966 | 322548 |
| MAPK_1.fcs | 10000 | 7871 | 6871 | 45 | 6826 | 10885 |
| MAPK_2.fcs | 10000 | 8097 | 7093 | 61 | 7032 | 16291 |
| MAPK_3.fcs | 10000 | 8328 | 7152 | 84 | 7068 | 19384 |
| MAPK_4.fcs | 10000 | 8403 | 7237 | 68 | 7169 | 15729 |
| Untransfected_BSA-FITC_1.fcs | 10000 | 8506 | 7569 | 80 | 7489 | 17316 |
| Untransfected_BSA-FITC_2.fcs | 10000 | 8617 | 7634 | 77 | 7557 | 15925 |
| Untransfected_BSA-FITC_3.fcs | 10000 | 8404 | 7339 | 116 | 7223 | 23611 |
| Untransfected_1.fcs | 10000 | 8407 | 7582 | 55 | 7527 | 10287 |
| Untransfected_2.fcs | 10000 | 8729 | 7991 | 23 | 7968 | 4750 |
| Untransfected_3.fcs | 10000 | 8371 | 7288 | 24 | 7264 | 5959 |

** As calculated from the sum of the intensities in the histogram of fluorescein + events*

**Table S4**

| Sample | total count | cell count | single cell count | Q1 count | Q2 count | Q3 count | Q4 count | BFP+ count | cells/Single Cells/BFP+ \| Mean (mScarlet-A) |
|---|---|---|---|---|---|---|---|---|---|
| BSA-F_1.fcs | 10000 | 8910 | 7161 | 213 | 682 | 570 | 5696 | 1252 | 22359 |
| BSA-F_2.fcs | 10000 | 8622 | 6842 | 227 | 686 | 479 | 5450 | 1165 | 22515 |
| MAPK_1.fcs | 10000 | 8855 | 7095 | 29 | 506 | 681 | 5879 | 1187 | 1191 |
| MAPK_2.fcs | 10000 | 8605 | 6831 | 30 | 271 | 357 | 6173 | 628 | 847 |
| MAPK_3.fcs | 10000 | 8658 | 6943 | 38 | 568 | 682 | 5655 | 1250 | 969 |
| no FITC_1.fcs | 10000 | 8620 | 6882 | 38 | 675 | 929 | 5240 | 1604 | 913 |
| no FITC_2.fcs | 10000 | 8152 | 6642 | 15 | 345 | 898 | 5384 | 1243 | 669 |
| no 3CP_1.fcs | 10000 | 8406 | 6559 | 56 | 453 | 644 | 5406 | 1097 | 3630 |
| no 3CP_2.fcs | 10000 | 8013 | 6133 | 69 | 632 | 771 | 4661 | 1403 | 4701 |
| untransfected_1.fcs | 10000 | 9194 | 7647 | 10 | 4 | 28 | 7605 | 32 | 76.4 |
| untransfected_2.fcs | 10000 | 9015 | 7497 | 10 | 5 | 23 | 7459 | 28 | 132 |
| untransfected_3.fcs | 10000 | 8644 | 6921 | 8 | 3 | 36 | 6874 | 39 | 80.3 |
| with 3CP_1.fcs | 10000 | 6403 | 5582 | 18 | 143 | 582 | 4844 | 725 | 650 |
| with 3CP_2.fcs | 10000 | 6452 | 5610 | 30 | 177 | 719 | 4684 | 896 | 650 |

### SUPPLEMENTARY INFORMATION ON MODEL DEVELOPMENT

##### 1. Derivation of the set of ordinary differential equations (ODEs)

We derive the mathematical model using ordinary differential equations (ODEs) in this section.

The model describes the binding of the receptor [Rec] to bovine serum albumin [BSA]

$$[\mathrm{Rec}] + [\mathrm{BSA}] \xrightarrow[k_{\mathrm{bind}}]{} [\mathrm{Rec\_BSA}], \tag{1.1}$$

which happens with the rate $k_{\mathrm{bind}}$. The bound complex can again bind another receptor

$$[\mathrm{Rec}] + [\mathrm{Rec\_BSA}] \xrightarrow[k_{\mathrm{bind\_complex}}]{} [\mathrm{Rec\_Rec\_BSA}] \tag{1.2}$$

with the rate $k_{\mathrm{bind\_complex}}$. Having recruited two receptors, the complex becomes active and induces the expression of $[\mathrm{SEAP_{State}}]$

$$\emptyset \xrightarrow[k_{\mathrm{express}}\,[\mathrm{Rec\_Rec\_BSA}]]{} [\mathrm{SEAP_{State}}]. \tag{1.3}$$

The full set of ordinary differential equations for the five states thus are

$$\frac{d[\mathrm{Rec}](t)}{dt} = -\,k_{\mathrm{bind}}\,[\mathrm{Rec}][\mathrm{BSA}] - k_{\mathrm{bind\_complex}}\,[\mathrm{Rec}]\,[\mathrm{Rec\_BSA}] \tag{1.4}$$

$$\frac{d[\mathrm{BSA}](t)}{dt} = -\,k_{\mathrm{bind}}\,[\mathrm{Rec}][\mathrm{BSA}] \tag{1.5}$$

$$\frac{d[\mathrm{Rec\_BSA}](t)}{dt} = +\,k_{\mathrm{bind}}\,[\mathrm{Rec}][\mathrm{BSA}] - k_{\mathrm{bind\_complex}}\,[\mathrm{Rec}]\,[\mathrm{Rec\_BSA}] \tag{1.6}$$

$$\frac{d[\mathrm{Rec\_Rec\_BSA}](t)}{dt} = +\,k_{\mathrm{bind\_complex}}\,[\mathrm{Rec}]\,[\mathrm{Rec\_BSA}] \tag{1.7}$$

$$\frac{d[\mathrm{SEAP_{State}}](t)}{dt} = +\,k_{\mathrm{express}}\,[\mathrm{Rec\_Rec\_BSA}] \tag{1.8}$$

#### 2. Implementation of the experiment

The data used to calibrate the model and determine its uncertainties is the dose-response measurement of SEAP activity for different BSA-fluorescein concentrations, shown in Figure 2c in the main text. The modelling starts at time point 0 h with the addition of medium containing BSA-fluorescein. SEAP activity is measured 20 h after addition of BSA.

The observation function of SEAP is

$$\frac{d[\mathrm{SEAP}](t)}{dt} = \mathrm{scale}_{\mathrm{SEAP}}\,[\mathrm{SEAP}_{\mathrm{State}}] + \mathrm{offset}_{\mathrm{SEAP}}. \tag{2.1}$$

Because the scaling of SEAP is unknown, the scaling parameter cannot be estimated from the data. Instead, it has to be fixed. We choose to fix the scaling to the maximal data value of the dataset (39.84 U/L), to increase the numerical efficiency of the model calculations. The offset of the SEAP measurements corresponds to the SEAP activity for a concentration of zero BSA-fluorescein. SEAP activity at this concentration was measured and the offset fixed to the mean of the six measured datapoints

$$\mathrm{offset}_{\mathrm{SEAP}} = 2.53\ [\mathrm{U/L}]. \tag{2.1}$$

The experimental error of the measurements was determined by calculating the variance of the replicates for the different concentrations and averaging over them. The standard deviation of the data was thus determined to be 2.37 [U/L].

#### 3. Initial conditions

The initial conditions of the model, i.e. the concentration of the five model states at time point zero are necessary to solve the model equations. No BSA-fluorescein-induced SEAP is present before the addition of BSA-fluorescein to the system, thus

$$[\mathrm{SEAP}_{\mathrm{State}}](0) = 0\ [\mathrm{a.u.}]. \tag{3.1}$$

Furthermore, at time point zero, the complexes

$$[\mathrm{Rec\_BSA}](0) = 0\ [\mathrm{nM}] \quad \text{and} \tag{3.2}$$

$$[\mathrm{Rec\_Rec\_BSA}](0) = 0\ [\mathrm{nM}] \tag{3.3}$$

are not yet formed.

The concentration of the receptor will be estimated as free parameter

$$[\mathrm{Rec}](0) = \mathrm{init}_{\mathrm{Rec}}[\mathrm{nM}], \tag{3.4}$$

since it is unknown.

Finally, the concentration of BSA is given through the experimental setup, i.e.

$$[\mathrm{BSA}](0) = \mathrm{input}_{\mathrm{concentration}}, \tag{3.5}$$

where $\mathrm{input}_{\mathrm{concentration}}$ corresponds to the experimentally added concentration of BSA in nM.

#### 4. Simplification of the model by solving practical non-identifiability

Using a maximum likelihood approach, we determined the unknown parameters of the model by comparing its predictions to the measured data.[1] For this purpose we performed a multi-start optimization with 100 runs starting from different, randomly drawn, initial parameter guesses. This yielded the maximum likelihood estimate of the parameters. The parameter optimization was performed on a logarithmic scale, accounting for the non-negativity of the rate constants.[3] Starting from the maximum likelihood estimate, we furthermore calculated the parameter profile likelihood of the unknown parameters to determine their confidence intervals.[4] The entire modelling process was performed using the Data2Dynamics software package freely available on GitHub.[5] The code and data of the modelling process are provided as supporting files with this publication to enable the reproduction of the modelling results.

The model was fitted to 24 data points and three dynamic parameters ($k_{\mathrm{bind}}$, $k_{\mathrm{bind\_complex}}$, and $k_{\mathrm{express}}$) as well as one unknown initial parameter ($\mathrm{init}_{\mathrm{Rec}}$) were estimated. The parameter profile of $k_{\mathrm{bind\_complex}}$ shown in Figure S5 shows, that this parameter is practically non-identifiable[4]. Since the likelihood is flat for large values of $k_{\mathrm{bind\_complex}}$, the parameter can be fixed[6]

$$k_{\mathrm{bind\_complex}} = 10^3 \; [\, 1 \,/\, (\mathrm{nM\ h}) \,]. \tag{4.1}$$

This reduces the model to two dynamical parameters and one unknown initial value.

[1] See the supporting information of Beyer et al.[2] for a detailed description of the modelling process (https://onlinelibrary.wiley.com/action/downloadSupplement?doi=10.1002%2Fadma.201800472&file=adma201800472-sup-0001-S1.pdf).

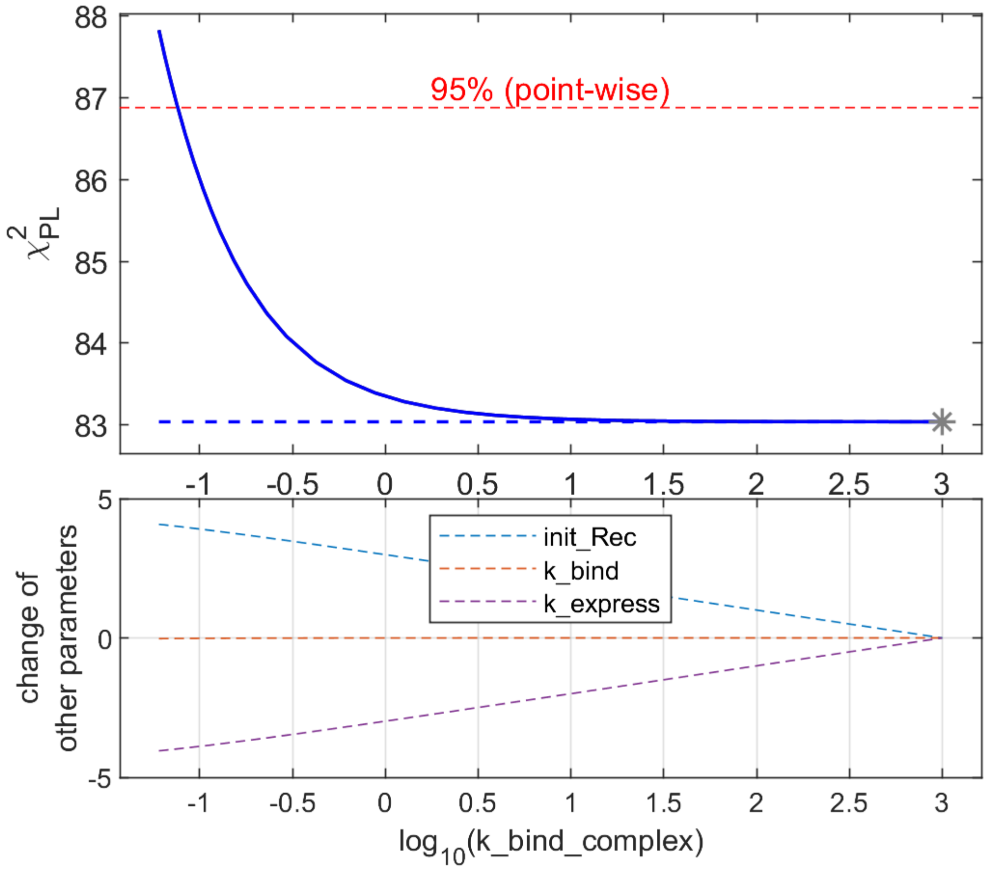


**Figure S5. Parameter profile likelihood of $k_{\text{bind\_complex}}$.** Parameter profile likelihood of the parameter $k_{\text{bind\_complex}}$ (above) and the changes of three other parameters along the profile likelihood of $k_{\text{bind\_complex}}$ (below). The parameter is practically non-identifiable at 95% point-wise confidence level (red dashed line), shown by the flat profile likelihood (blue line) to the right converging to the likelihood value of the maximum likelihood estimate (blue dashed line). The grey asterisk signifies the maximum likelihood estimate of the model. In the lower graph, the change of parameters of the three other parameters from the maximum likelihood estimate is shown. The parameter changes of the other parameters show the linear dependence of $\text{init}_{\text{Rec}}$ and $k_{\text{express}}$ on $k_{\text{bind\_complex}}$.

#### 5. Results of the parameter estimation and uncertainty analysis

After fixing $k_{\text{bind\_complex}}$, the multi-start optimization was again performed with 100 runs. The model was thus again fitted to 24 data points and now two dynamic parameters as well as one unknown initial parameter were estimated. The results of the multi-start optimization showed that 95 runs converged to the lowest optimum, suggesting it is the global optimum (Figure S6).

The parameter profile likelihood of the three parameter showed that all three are identifiable and have finite confidence intervals (Figure S7). Their numerical values and the detailed results of the parameter estimation are shown in Table S5.

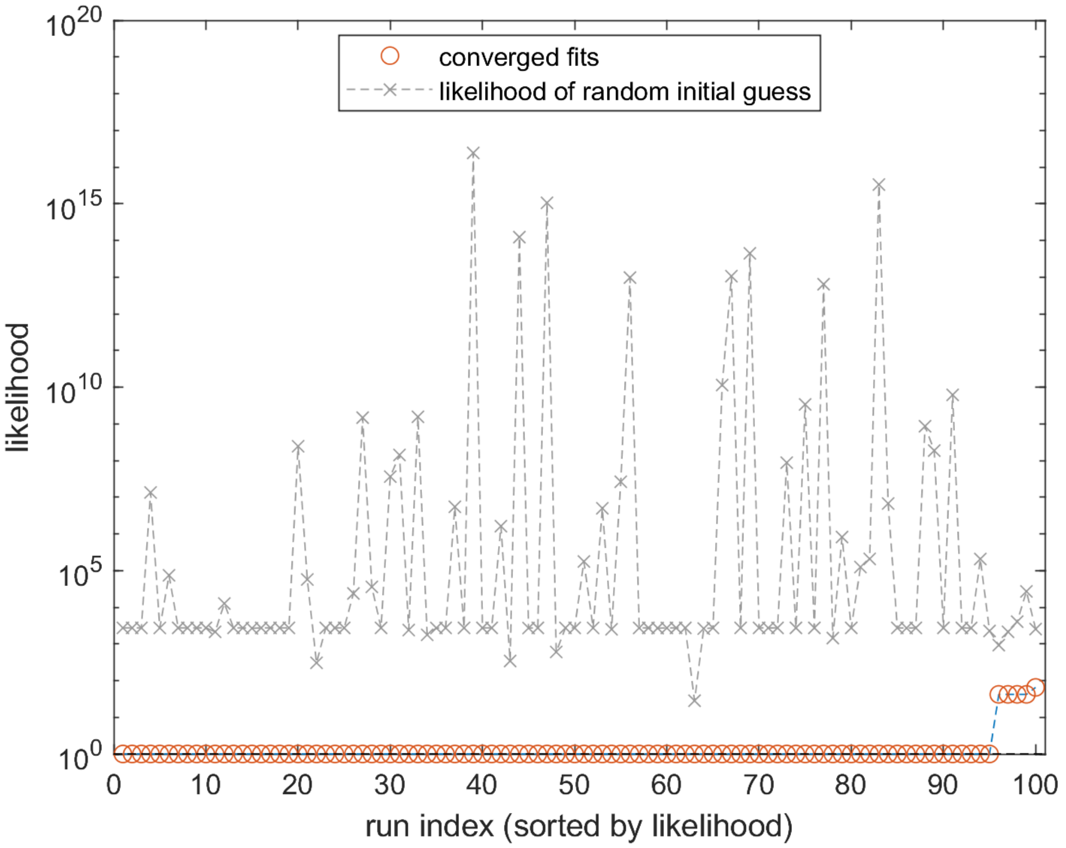


**Figure S6. Multi-start optimization results**. Result of 100 optimization runs with random initial parameter guesses. The y-axis shows the likelihood value, normalized to the lowest value. 95 runs converged to the lowest likelihood value, suggesting it is the global optimum.

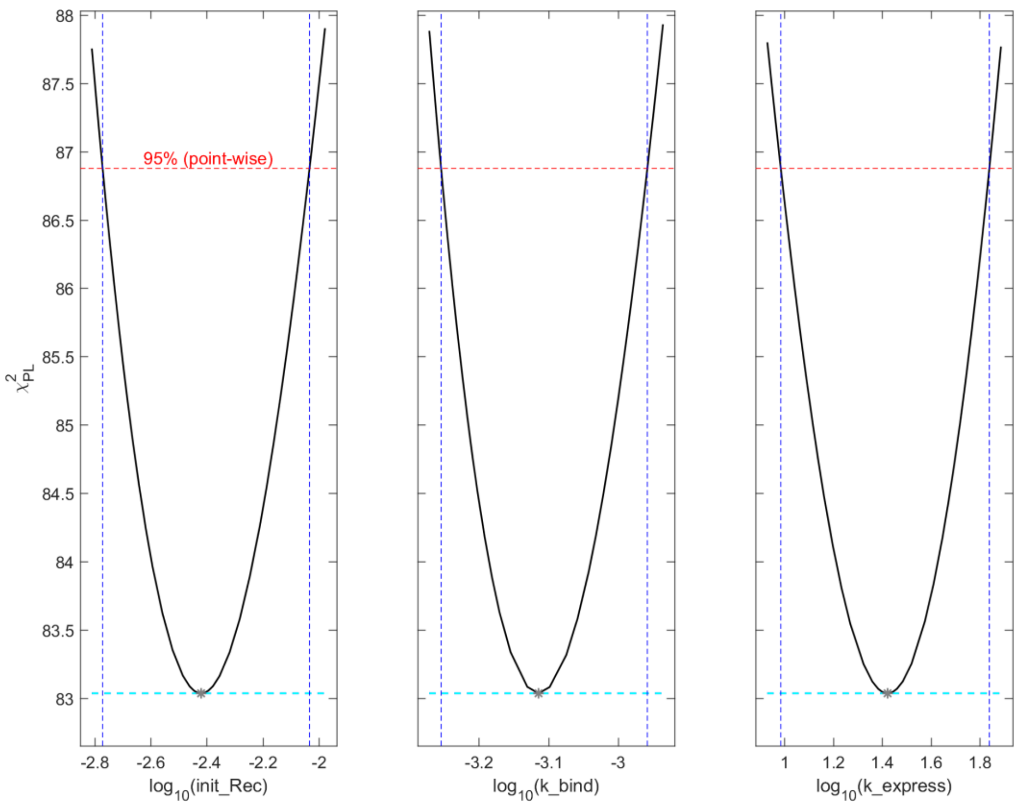


**Figure S7. Parameter profile likelihood**. Parameter profile likelihood of the three free parameters of the model. All parameters are identifiable with finite 95% point-wise confidence intervals, signified by the intersection of the parameter profiles (black line) with the 95% point-wise confidence level (red dashed line). This intersection is visualized through the blue dashed line. The turquoise dashed line signifies the likelihood value of the maximum likelihood estimate of the model. The grey asterisk shows the maximum likelihood estimate. The parameter axis shows the parameter value on logarithmic scale.

**Table S5. Estimated model parameters**. The three estimated model parameters on natural scale, their maximum likelihood estimate and 95% point-wise confidence intervals as given by the profile likelihood method.

| Parameter | Maximum likelihood estimate | Confidence intervals |
|---|---|---|
| $\mathbf{init_{Rec}}$ | 0.0038 [nM] | [0.0017 , 0.0092] |
| $\boldsymbol{k}_{\mathbf{bind}}$ | 0.00077 [1/(nM h)] | [0.00060 , 0.00110] |
| $\boldsymbol{k}_{\mathbf{express}}$ | 26 [1/(nM h)] | [9.6 , 68.8] |
